# A Systematic Review of the Neuroprotective Role and Biomarker Potential of GDF15 in Neurodegeneration

**DOI:** 10.1101/2024.07.07.600156

**Authors:** Finula I. Isik, Shannon Thomson, John F. Cueto, Jessica Spathos, Samuel N. Breit, Vicky W.W. Tsai, David A. Brown, Caitlin A. Finney

## Abstract

Neurodegeneration is characteristically multifaceted, with limited therapeutic options. One of the chief pathophysiological mechanisms driving these conditions is neuroinflammation, prompting increasing clinical interest in immunomodulatory agents. Growth differentiation factor 15 (GDF15; previously also called macrophage inhibitory cytokine-1 or MIC-1), an anti-inflammatory cytokine with established neurotrophic properties, has emerged as a promising therapeutic agent in recent decades. However, methodological challenges and the delayed identification of its specific receptor GFRAL have hindered research progress. This review systematically examines literature about GDF15 in neurodegenerative diseases and neurotrauma. The evidence collated in this review indicates that GDF15 expression is upregulated in response to neurodegenerative pathophysiology and increasing its levels in preclinical models typically improves outcomes. Key knowledge gaps are addressed for future investigations to foster a more comprehensive understanding of the neuroprotective effects elicited by GDF15.

## 1. Introduction

Neurodegenerative disorders, characterised by the gradual loss of neurons, have limited treatment options and present a significant diagnostic challenge in modern healthcare. These conditions result from a range of chronic diseases driven by complex neuropathogenic pathways or secondary to neurotrauma. Neuroinflammation is intimately associated with these events and can both mitigate and potentiate ongoing neuronal damage. On one hand, inflammatory mechanisms are integral in removing cellular debris and promoting repair and regeneration. On the other, chronic, inappropriately amplified, or suppressed responses, can exacerbate damage or propagate ongoing inflammatory cascades involved in generating neuropathology. Neuroinflammatory sequelae are now widely acknowledged to have an important role in responding to and/or promoting endogenous, neurodegenerative disease-specific protein aggregates including, but not limited to, α-synuclein, amyloid-β, hyperphosphorylated tau and more (reviewed in Patani et al., 2023, Passaro et al., 2021, Skaper et al., 2018). Therefore, immunomodulation represents a promising avenue for the treatment of neurodegenerative conditions and are the basis of ongoing preclinical and clinical research. To date, however, most anti-inflammatory drugs fail to demonstrate significant, favourable therapeutic effects (Scott et al., 2017). These findings may be due to the suppression of key, beneficial inflammatory events, highlighting that ‘pro-inflammatory’ events at appropriate times may have a reparative rather than damaging immunomodulatory phenotype (Scott et al., 2017). Therefore, there is a pressing need to characterise the involvement of key inflammatory molecules in neurodegeneration and to identify novel therapeutic targets that can mitigate or prevent neurodegenerative processes in the central nervous system (CNS).

Growth differentiation factor 15 (GDF15; previously called macrophage inhibitory cytokine-1 MIC-1) is one such immunomodulatory factor that has been implicated in neurodegeneration. This GDNF family cytokine, within the transforming growth factor-β (TGF-β) superfamily, was first identified as a 25kDa disulfide-linked dimer (Bootcov et al., 1997). Like other TGF-β superfamily members, it is synthesized as a precursor protein that dimerises and is processed by cleavage at the conserved RXXR sequence, resulting in the secretion of a mature, biologically active disulfide-linked dimer that rapidly diffuses into circulation. Unprocessed protein is sometimes secreted and binds to the extracellular matrix, possibly serving as a local reservoir (Bauskin et al., 2005). Centrally, GDF15 regulates non-homeostatic energy metabolism through its only known receptor, growth factor receptor α-like (GFRAL), expressed on neurons in the hindbrain area postrema (AP) and nucleus of the solitary tract (NTS). GDF15 and GFRAL form a complex with the <u>RE</u>arranged during <u>T</u>ransfection (RET) coreceptor to activate these hindbrain neurons (Hsu et al., 2017, Mullican et al., 2017, Emmerson et al., 2017, Yang et al., 2017, Breit et al., 2021). Until recently, GDF15 was incorrectly believed to act via classical TGF-β receptors I and II and SMAD pathways. Many *in vitro* studies demonstrating this effect used commercial recombinant GDF15 (rGDF15) stocks which are now known to be contaminated with TGF-β, rendering many conclusions related to the GDF15’s effects unreliable (Olsen et al., 2017). Regardless, accumulating work utilising transgenic mouse models indicate GDF15 can modulate peripheral immune cell infiltration and contribute to CNS/cardiac infarct or lesion healing (Preusch et al., 2013, de Jager et al., 2011, Golakani et al., 2019, Kempf et al., 2011, Husaini et al., 2020). Moreover, this pleiotropic molecule is a well-established indicator of cellular stress. The GDF15 promotor has binding sites for many transcriptional regulators induced by cell stress, including p53 (Li et al., 2000), EGR-1 (Baek et al., 2004), CHOP (Chung et al., 2017) and ATF4 (Patel et al., 2019). Elevated serum GDF15 is a feature of pregnancy and frequently observed in conditions like advanced cancers, chronic heart and renal failure, genetic mitochondrial diseases, obesity and type 2 diabetes, dementia and chronic inflammatory diseases (Breit et al., 2021), and is a reliable predictor of all-cause mortality (Moon et al., 2020, Wiklund et al., 2010). Despite the first evidence of GDF15’s neuroprotective role reported over two decades ago (Subramaniam et al., 2003), its role as a central or peripheral neurodegenerative biomarker and its potential neuroprotective effects remain poorly defined.

To address this gap in the literature and begin to elucidate the role of GDF15 in neurodegeneration, we performed a systematic review of studies examining this pleiotropic cytokine in neurodegenerative conditions. Key findings drawn from these studies were categorised into one of two main themes, which will be outlined in this present review: (1) investigating the neuroprotective role of GDF15 in animal and cell models of neurodegeneration, and (2), assessing GDF15 levels as a biomarker for neurodegenerative disease or injury. In doing so, we provide the first systematic examination of literature on GDF15 in relation to neurodegenerative physiopathology and highlight the potential for GDF15 as a viable therapeutic target across several neurodegenerative diseases.

## 2. Methods

### 2.1. Literature search

For the present systematic review, we employed a clear pipeline as previously detailed (Shvetcov et al., 2023). In brief, we systematically searched three separate databases: PubMed, Scopus, and Web of Science. Our search terms included GDF15 and its synonyms (e.g. MIC-1) as well as terms related to the CNS (e.g. brain), as detailed in Table 1.

**Table 1.**
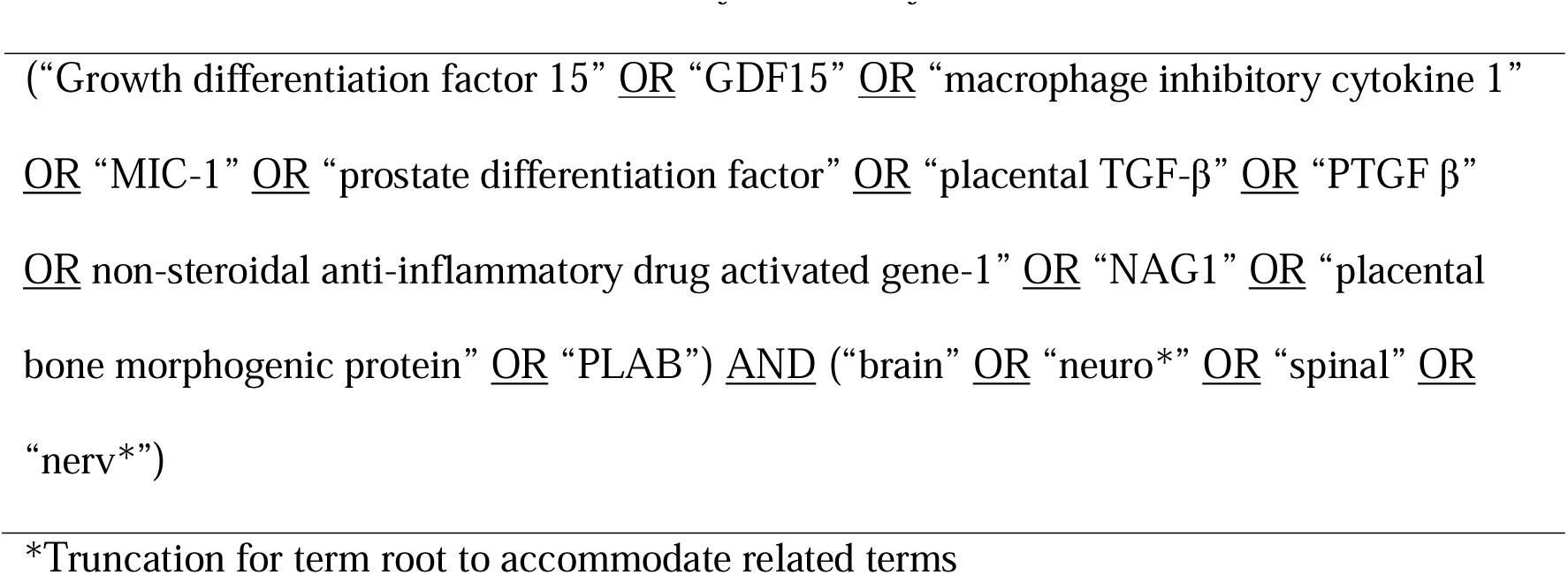
Key terms for database search. The search strategy incorporated multiple known names for GDF15 in combination with key nervous system terms.

The search terms were limited to title and abstract only and databases were searched on May 27, 2024. Following removal of duplicates, titles, abstracts, and full texts were reviewed and scored for inclusion by F.I. and S.T. based on eligibility criteria developed *a priori* (Table 2).

**Table 2.**
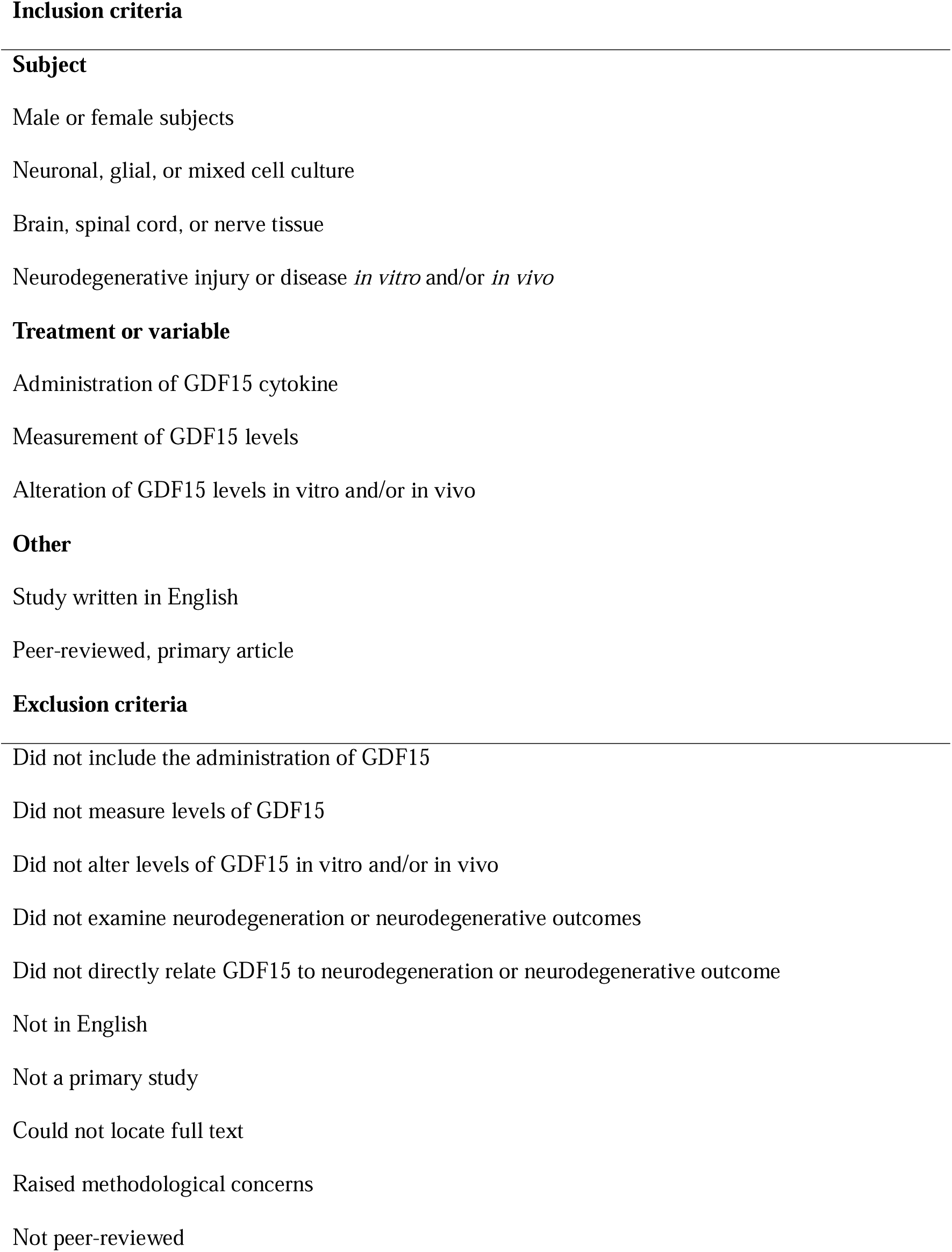
Inclusion and exclusion selection criteria.

### 2.2. Eligibility criteria

Our goals in this systematic review were to assess GDF15’s involvement in neurodegenerative conditions to (1) elucidate the mechanisms that may underly its neuroprotective effects and (2) assess its suitability as a biomarker for neurodegenerative diseases. We therefore developed clear inclusion and exclusion criteria to address this (Table 2). Inclusion criteria included (1) a human, animal or cell model of neurodegenerative injury or insult, (2) confirmed neurodegenerative injury (*in vitro* or *in vivo*), (3) exogenous manipulation of GDF15 or direct measurement of GDF15, (4) study written in English and (5) a peer reviewed primary article. The studies were excluded if they did not fit these inclusion criteria, namely, they (1) did not include the measurement of, alteration of, or exogenous modulation of GDF15 levels *in vitro* and/or *in vivo*, (2) did not examine neurodegeneration or neurodegenerative outcomes, (3) did not directly relate GDF15 to neurodegeneration or neurodegenerative outcomes, (4) were not in English, (5) were not peer reviewed, (6) not a primary study, and/or (7) were limited by methodological concerns, including inadequate or inconsistent result reporting. We took a liberal approach to screening titles to prevent the inappropriate exclusion of full texts. Two assessors (F.I. and S.T.) were used in the screening of titles and abstracts, and wherever a discordance was identified, the citation defaulted to having its full text assessed.

## 3. Results

### 3.1. Literature search

The literature search conducted in PubMed, Scopus and Web of Science resulted in a total 1,521 citations. After removing duplicate citations, 717 remained. These citation titles and abstracts were then screened based on the eligibility criteria (Table 2). The interrater reliability indicated substantial agreement between both raters (91.77%, Cohen’s Kappa: 0.71). A total of 153 full texts were then reviewed for inclusion. Of these, 83 full texts met the eligibility criteria and were included in the review (Figure 1). The most common reason for exclusion was not examining neurodegeneration or neurodegenerative outcomes (n = 41). Publications were additionally excluded if they did not examine the relationship between neurodegenerative variables and GDF15 (n = 11). Other reasons for exclusion included studies that did not include the measurement, modulation, or exogenous application of GDF15 *in vivo* or *in vitro* (n = 3), were not in English (n = 2), were not a primary study (n = 5) and texts that could not be located despite our best efforts (n = 4). Three papers were excluded due to concerns related to inconsistencies in results reported in graphs, questioning the validity of the reported findings.

**Figure 1.**
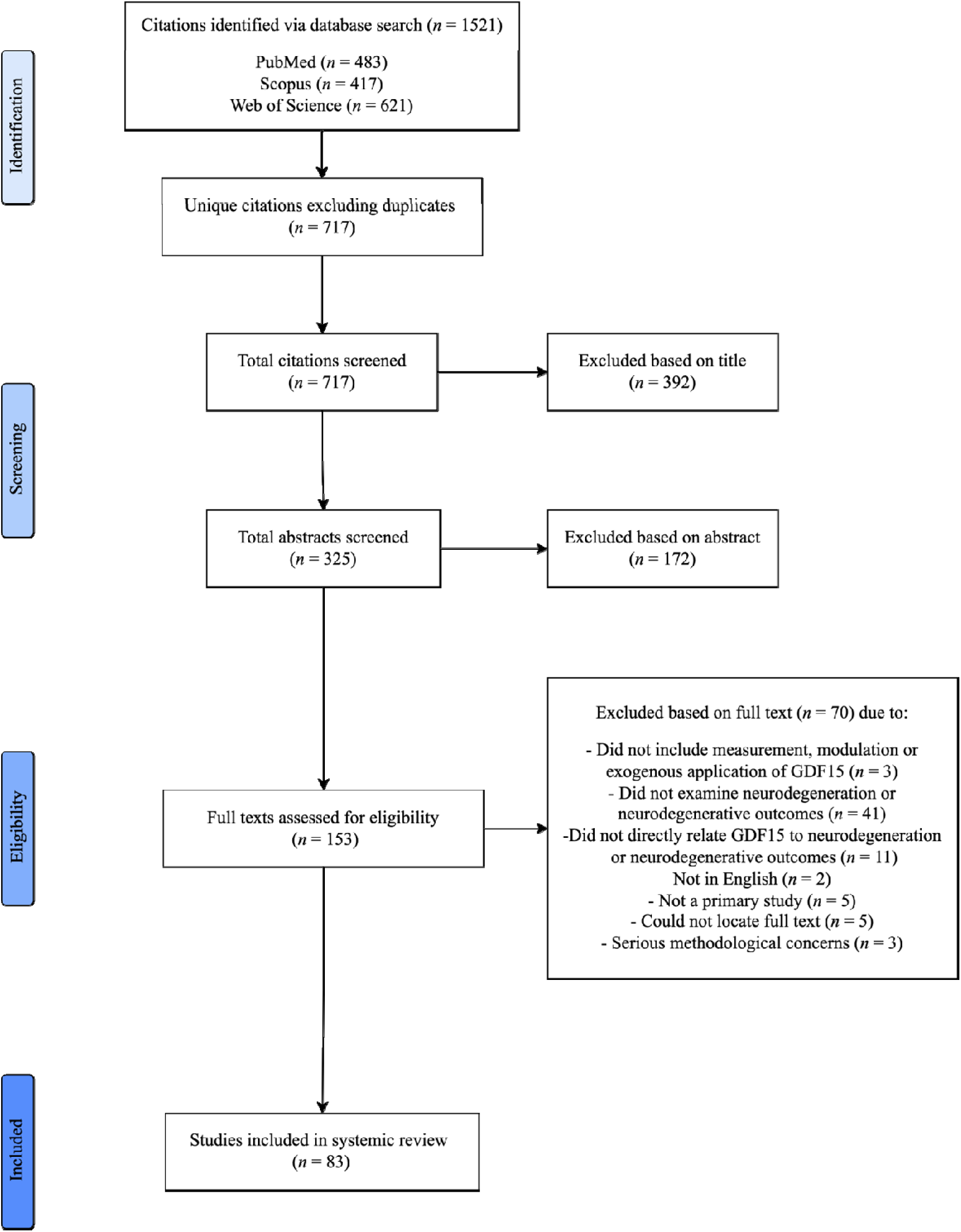
Systematic literature review process and result pipeline. The stages of literature identification, screening, exclusion, and inclusion to final full text.

### 3.2. Characteristics of the included studies

The included studies examined GDF15 in humans (n = 52), animal models (n = 22), or cell culture models (n = 26). Several studies included a combination of subjects, including animal and cell (n = 11), human and cell (n = 3) and human and animal (n = 2), and all three subjects (n = 1).

In human studies (Table 3), GDF15 levels were measured in biological fluids (n = 50) and/or post-mortem CNS tissue (n = 3) of individuals affected by neurodegenerative disease or injury. This included mitochondrial disease (n = 14), Alzheimer’s disease (AD) or dementia (n = 13), synucleinopathy (n = 7), Multiple Sclerosis (MS) and/or Neuromyelitis Optica spectrum disorder (n = 7), glaucoma (n = 3), motor neuron diseases (MND) (n = 3), Charcot-Marie-Tooth disease (n = 2), Vanishing White Matter Disease (n = 1), stroke (n = 8) and subarachnoid haemorrhage (n = 1). Most studies (n = 33) involved cross-sectional analysis of GDF15 levels in blood, cerebrospinal fluid (CSF) or aqueous humour. The remaining studies (n = 18) were longitudinal. They either measured GDF15 levels across several timepoints, or related baseline GDF15 levels with follow-up neurological outcomes. One case study and one 2-sample Mendelian Randomization Study were also included. Full characteristics related to study design and participants are outlined below and in Supplementary Tables 1 and 2.

**Table 3.** Characteristics of human subjects.

| **Human Subject Characteristics** | **Number of studies featuring cohort** |
| --- | --- |
| **Disease** |  |
| Mitochondrial disease | 14 |
| Alzheimer's disease (AD) and dementia | 13 |
| Synucleinopathy | 7 |
| Multiple Sclerosis (MS) /Neuromyelitis Optica Spectrum disorder (NMOSD) | 7 |
| Glaucoma | 3 |
| Motor neuron disease (MND) | 3 |
| Charcot-Marie-Tooth disease (CMT) | 2 |
| Vanishing White Matter disease (VWMD) | 1 |
| Stroke | 8 |
| Subarachnoid haemorrhage | 1 |
| **GDF15 Measurement** |  |
| Serum | 26 |
| Plasma | 18 |
| Blood | 2 |
| CSF | 3 |
| Aqueous humour | 3 |
| Ex vivo tissue | 3 |

For animal studies (Table 4), most used mice (n = 19), followed by rats (n = 5) and drosophila (n = 1). Several diseases were modelled, including AD (n = 3), Parkinson’s disease (PD) (n = 3), Charcot-Marie Tooth disease (n = 2), retinal degeneration and glaucoma (n = 3), Huntington’s disease, Spinal Muscular Atrophy, and Vanishing White Matter disease (n = 1, respectively). Studies also examined neurodegeneration secondary to induced CNS (n = 5) and peripheral NS (PNS) injury (n = 2), including spinal cord injury (SCI), cryolesion, excitotoxicity, stroke, and sciatic nerve crush. Cell culture studies (Table 5) used models derived from human (n = 16), mouse (n = 6), and rat (n = 10) origin. Most used neurons (n = 14), followed by astrocytes (n = 6), mixed midbrain cultures (n = 4), microglia (n = 3), brain-derived endothelia (n =1), retinal ganglion cells (n = 1) and Schwann cells (n =1). Patient derived fibroblasts (n = 2), mesenchymal stem cells (n = 2), neural stem cells (n = 1) and patient derived mesenchymal stromal cells (n = 1) were also used. Like animal studies, neurodegenerative disease was induced in these cells to model AD (n = 4), PD (n = 5), Amyotrophic Lateral Sclerosis (ALS), Charcot-Marie Tooth disease, Huntington’s disease, MS, retinal degeneration, Spinal Muscular Atrophy and Vanishing White Matter disease (n = 1, respectively). The remaining studies induced cellular injury (n = 9), via neuroinflammation, low potassium, ferroptosis, excitotoxicity, mitochondrial dysfunction, or DNA damage. GDF15 secretion and expression was measured in cell and animal models of neurodegeneration (n = 26). Full characteristics of *in vitro* and *in vivo* studies are outlined in Supplementary Table 3.

**Table 4.** Characteristics of animal subjects.

| **Animal Characteristics** | **Number of studies featuring animal** |
| --- | --- |
| **Species, strain** |  |
| Mouse, all | 19 |
| Rat, all | 5 |
| Drosophila, HTT93Q exon1 | 1 |
| **Neurodegenerative Disease/ Condition Model** |  |
| AD | 3 |
| Parkinson's disease (PD) | 3 |
| Retinal degeneration (RD) /Glaucoma | 4 |
| Huntington's disease (HD) | 1 |
| Spinal Muscular Atrophy Jokela (SMAj) | 1 |
| Vanishing White Matter disease (VWMD) | 1 |
| CNS Injury | 5 |
| PNS Injury | 2 |

**Table 5.**
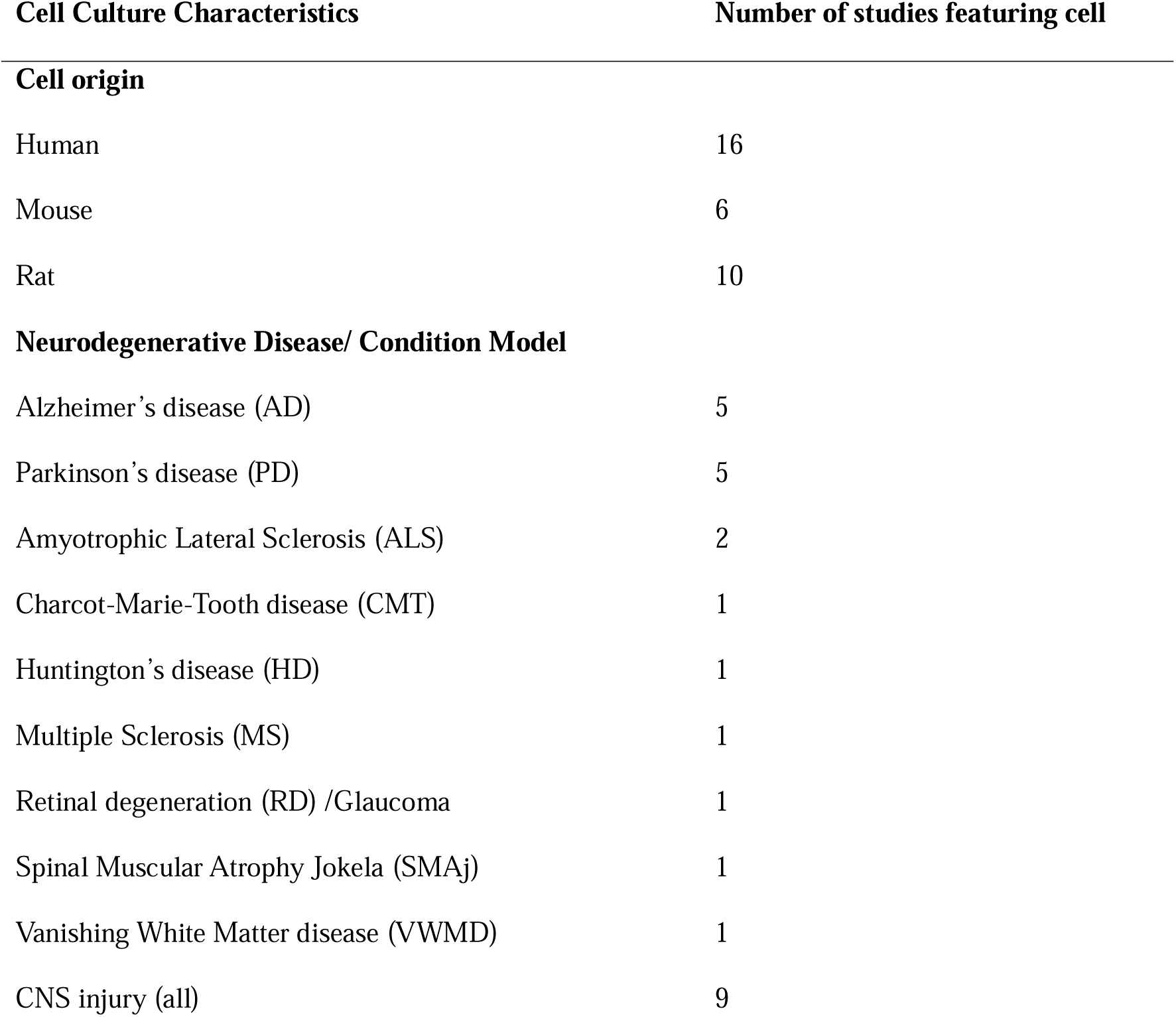
Characteristics of Cell Culture subjects.

| **Cell Culture Characteristics** | **Number of studies featuring cell** |
| --- | --- |
| **Cell origin** |  |
| Human | 16 |
| Mouse | 6 |
| Rat | 10 |
| **Neurodegenerative Disease/ Condition Model** |  |
| Alzheimer's disease (AD) | 5 |
| Parkinson's disease (PD) | 5 |
| Amyotrophic Lateral Sclerosis (ALS) | 2 |
| Charcot-Marie-Tooth disease (CMT) | 1 |
| Huntington's disease (HD) | 1 |
| Multiple Sclerosis (MS) | 1 |
| Retinal degeneration (RD) /Glaucoma | 1 |
| Spinal Muscular Atrophy Jokela (SMAj) | 1 |
| Vanishing White Matter disease (VWMD) | 1 |
| CNS injury (all) | 9 |

### 3.3. GDF15 is involved in neuroprotective molecular mechanisms in neurodegeneration

The animal and cell culture studies included in this systematic review highlight GDF15’s neuroprotective role across a wide range of pathophysiological mechanisms. In general, they showed that increasing GDF15 was beneficial, whilst decreasing its levels was detrimental. It should be noted that many of these studies used commercial rGDF15 protein, which is known to be a source of TGF-β contamination (Olsen et al., 2017). This makes it difficult to solely attribute these protective effects to our cytokine of interest. Although we have included details of these flagged studies (see Tables 6 – 8), we only discuss the results of studies that are not affected by this issue.

Overall, the studies examining GDF15 in neurodegenerative models highlight its role in minimising induced cell death (Table 6). Specifically, apoptosis was repeatedly demonstrated to be mitigated by GDF15 modulation. Increasing GDF15 levels *in vitro* decreased the proportion of PI positive (Subramaniam et al., 2003, Xiong et al., 2023) and TUNEL positive cells (Subramaniam et al., 2003, Li et al., 2022a), as well as reduced DNA fragmentation caused by apoptotic damage. Further, anti-apoptotic genes Bcl-2, PGCI-α and TH were increased whilst pro-apoptotic Bad, Bax, caspase-3 and P53 were decreased (Li et al., 2022a, Xiong et al., 2023). Reciprocal findings were demonstrated with knockdown of GDF15 in culture (Xiong et al., 2021) and GDF15 knockout mice (Charalambous et al., 2013). The anti-apoptotic effects of GDF15 were mediated by activating the PI3K/Akt pathway and blocking ERK activation. The protective effect of GDF15 on cerebellar granule cells exposed to low potassium was ablated by selective PI3K inhibitors LY294002 and wortmannin (Subramaniam et al., 2003). Additionally, low potassium induced ERK and in turn c-Jun activation but was prevented by GDF15 (Subramaniam et al., 2003). A more recent publication supported this by highlighting that the PI3K/Akt signalling pathway was significantly enriched in oligomycin treated neurons that overexpressed GDF15 (Liu et al., 2019). As GDF15 is known to phosphorylate Erk1/2 *in vivo* (Johnen et al., 2007), it is therefore likely that a neuroprotective effect of GDF15 occurs via this pathway.

**Table 6.**
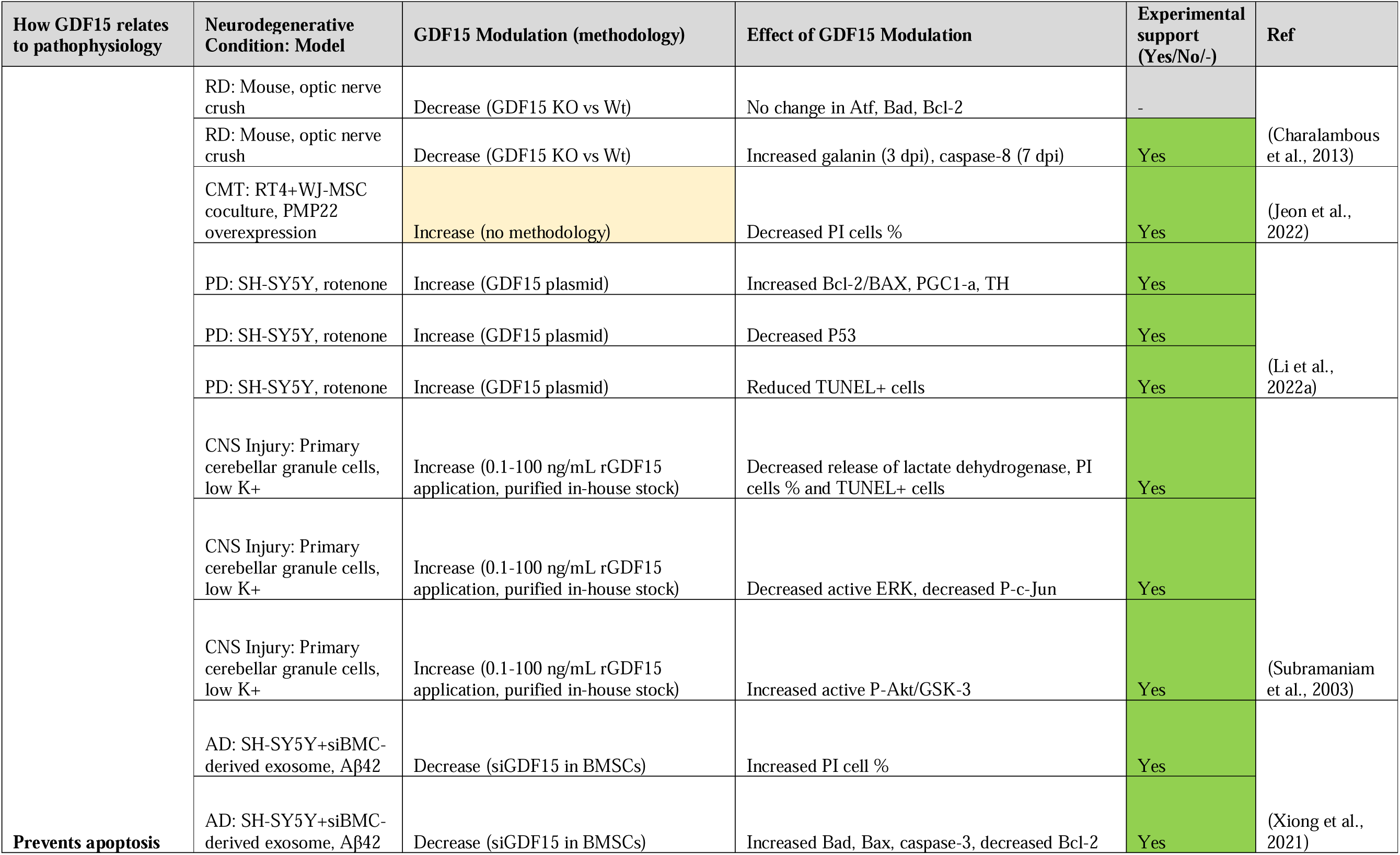
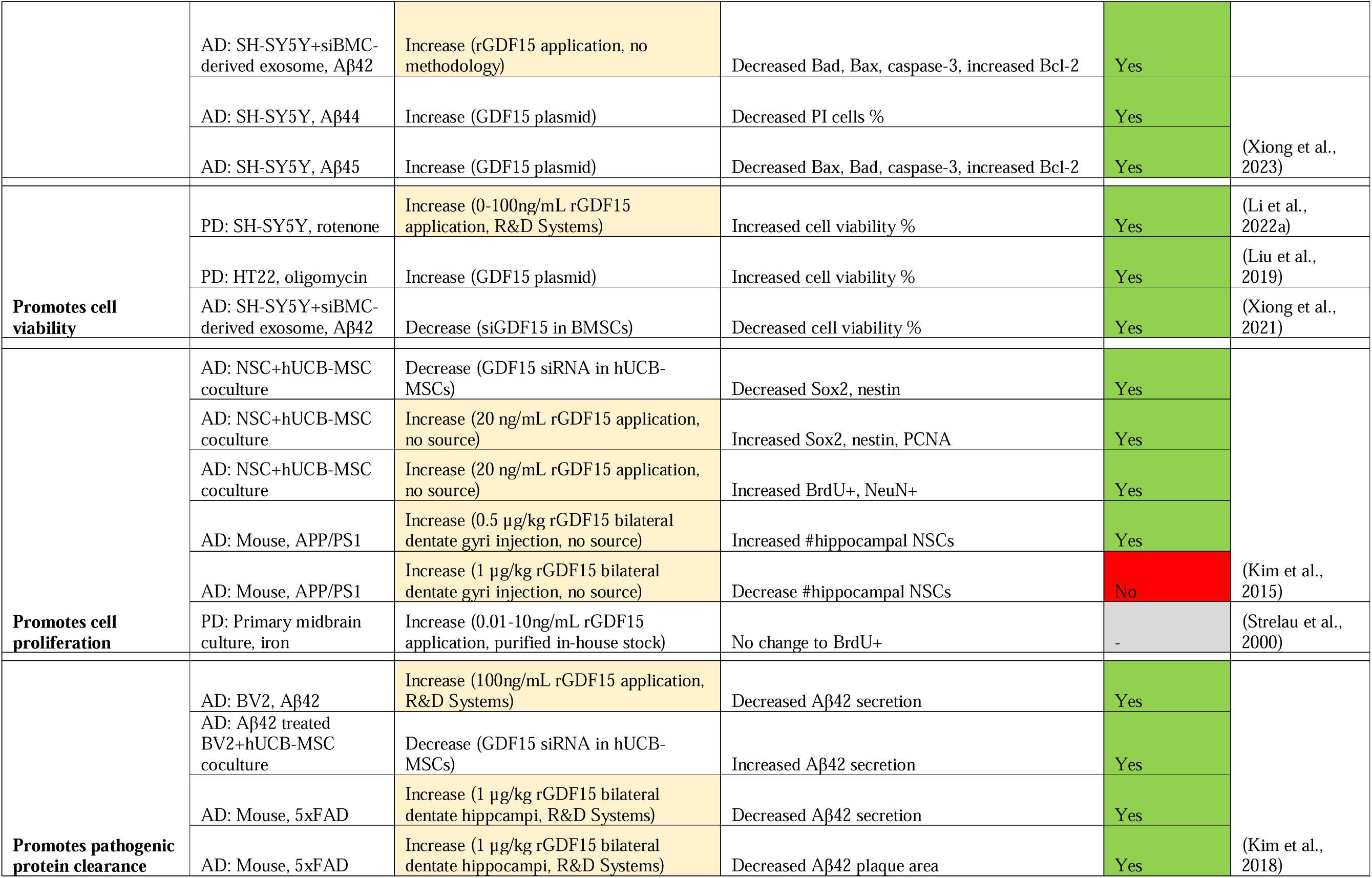
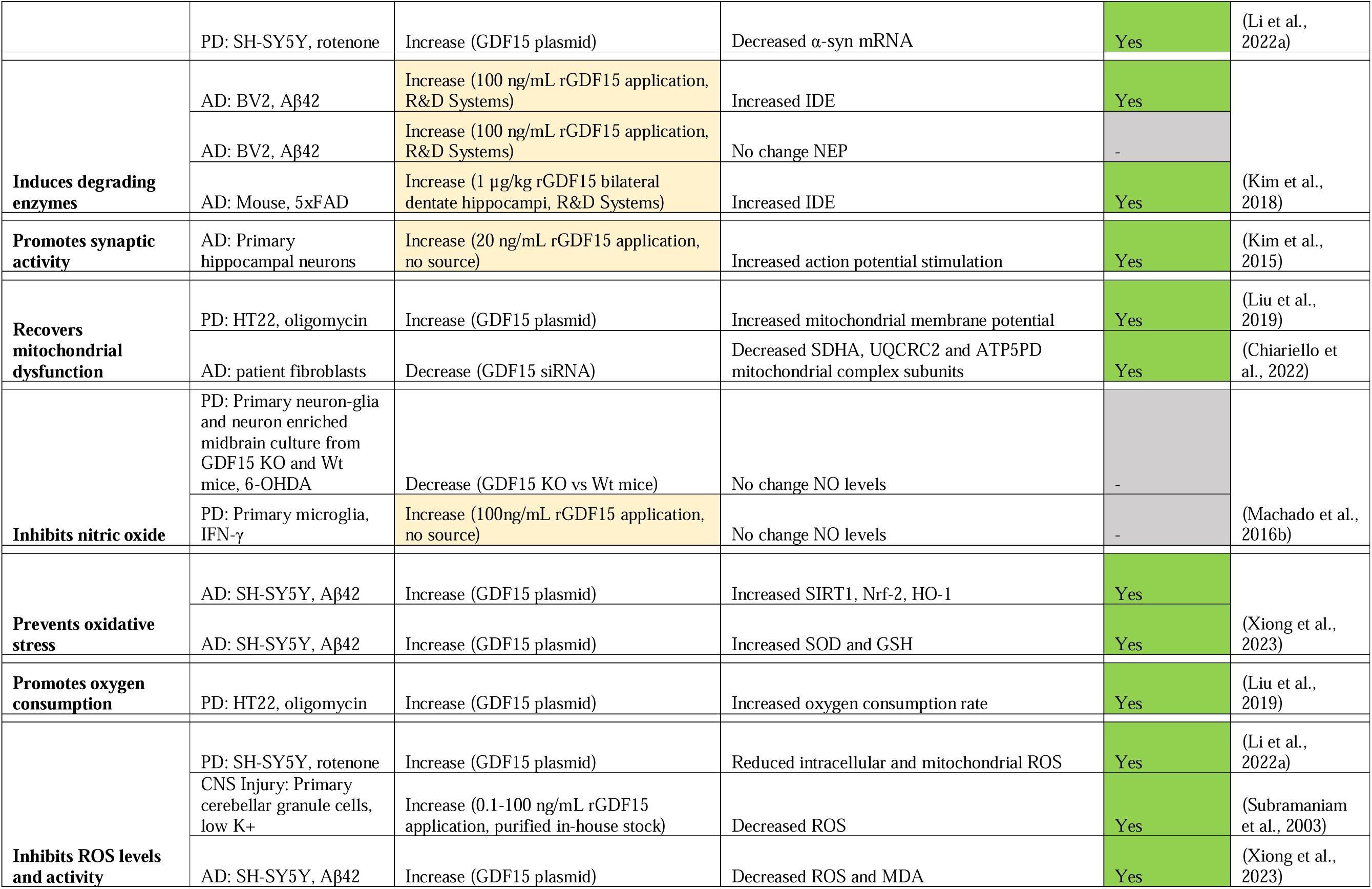
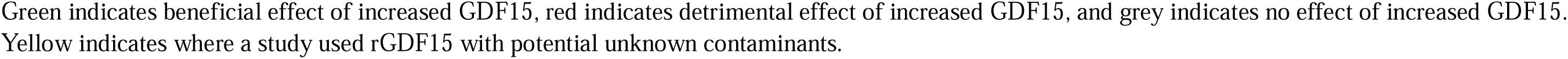
Modulating GDF15 in ND models is neuroprotective by promoting cell survival.

Previous reports highlight GDF15-GFRAL binding initiates RET signalling cascades including activation of Akt, Fos, Erk1/2 and phospholipase C (Breit et al., 2021), however, only three studies in this present review directly examined this receptor and its signalling cascade. GFRAL expression was identified in dopaminergic neurons (Kostuk et al., 2019), primary retinal ganglion cells (Iwata et al., 2021), and in mouse retina and optic nerve (Iwata et al., 2021), although, expression in these regions has not been identified in other publications. Transcriptomic analysis of wildtype and ALS SOD1^G93A^ mice failed to show GFRAL expression in spinal cord tissue (Younes et al., 2023). The neurotrophic effect of the GDF15-GFRAL-RET complex was demonstrated in retinal ganglion cells using RET inhibition. Specifically, rGDF15 activated Akt and promoted neurite and axon length in these cells. This effect was suppressed to the level of vehicle-control when co-treated with selective RET inhibitor GSK317910, which was also shown to suppress Akt phosphorylation (Iwata et al., 2021). Though this study used rGDF15 derived from *E. coli*, which may have residual endotoxin contamination or may have been misfolded, it provides intriguing support for further investigation. Further work utilising purified rGDF15 should be undertaken to confirm the mechanism of GDF15-GFRAL RET receptor-ligand interactions in Akt-mediated cell survival responses.

In addition to anti-apoptotic activity, GDF15 further promotes neuronal survival by increasing cell viability (Table 6). One study highlighted that genetically increasing GDF15 improved viability of HT22 cells in response to mitochondrial toxin oligomycin (Liu et al., 2019). Another study reinforced this by examining the effect of incubating cells with exosomes extracted from siGDF15 transfected mesenchymal stem cells (MSCs). When co-cultured with SH-SY5Y cells, these exosomes potentiated Aβ-induced loss of cell viability (Xiong et al., 2021). In addition, GDF15 was shown to regulate adult neural stem cell (NSC) proliferation. Normally, MSCs boost NSC proliferation through secretion of transcription factors Sox2 and nestin, an effect that is inhibited in siGDF15 MSC transfected co-cultures (Kim et al., 2015). However, a different study showed the application of rGDF15 failed to increase BrdU+ staining in iron-treated midbrain cultures (Strelau et al., 2000). Therefore, while GDF15 may be neurotrophic, there is insufficient evidence that supplementing exogenously can increase neuronal proliferation following injury.

GDF15 also has a positive impact on bioenergetic stress following injury and insult (Table 6). One study showed that reducing GDF15 in AD patient-derived fibroblasts led to a decrease in expression of mitochondrial complex subunits SDHA, UQCRC2 and ATP5PD (Chiariello et al., 2022). Other publications highlighted that increasing GDF15 levels *in vitro* helped restore mitochondrial function and reduced oxidative stress. Specifically, GDF15 plasmid transfection of neurons attenuated mitochondrial injury by restoring mitochondrial membrane potential and oxygen consumption (Liu et al., 2019), reducing ROS levels (Li et al., 2022a, Xiong et al., 2023, Subramaniam et al., 2003), and increasing expression of antioxidants SOD and GSH in response to Aβ42 (Xiong et al., 2023). This in turn promotes cell survival by inhibiting ROS-induced activation of pro-apoptotic ERK pathways (Subramaniam et al., 2003).

### 3.4. GDF15 may promote functional recovery by attenuating neuron loss and damage

Modulation of GDF15 in the included studies was partially associated with improved functional outcomes in neurodegenerative models (Table 7). An early study by Strelau and colleagues demonstrated functional motor recovery could be promoted in 6-OHDA treated rats via unilateral injection of rGDF15 into substantia nigra/lateral ventricle. Higher doses decreased rotational asymmetry, and low doses decreased amphetamine-induced rotations (Strelau et al., 2000). Complementing this, GDF15 knockout mice were shown to have poorer rotarod performance (Strelau et al., 2009). GDF15 overexpressing mice with SCI have improved motor scores from 7 days post injury (dpi) onwards, although the GDF15 null mice did not differ from the wildtype (Golakani et al., 2019). Similarly, GDF15 knockout genotype did not influence electromyography measures of nerve conductivity after sciatic nerve lesion (Wang et al., 2015), highlighting that while increased GDF15 improved motor performance following injury, a lack of GDF15 did not necessarily impede recovery.

**Table 7.**
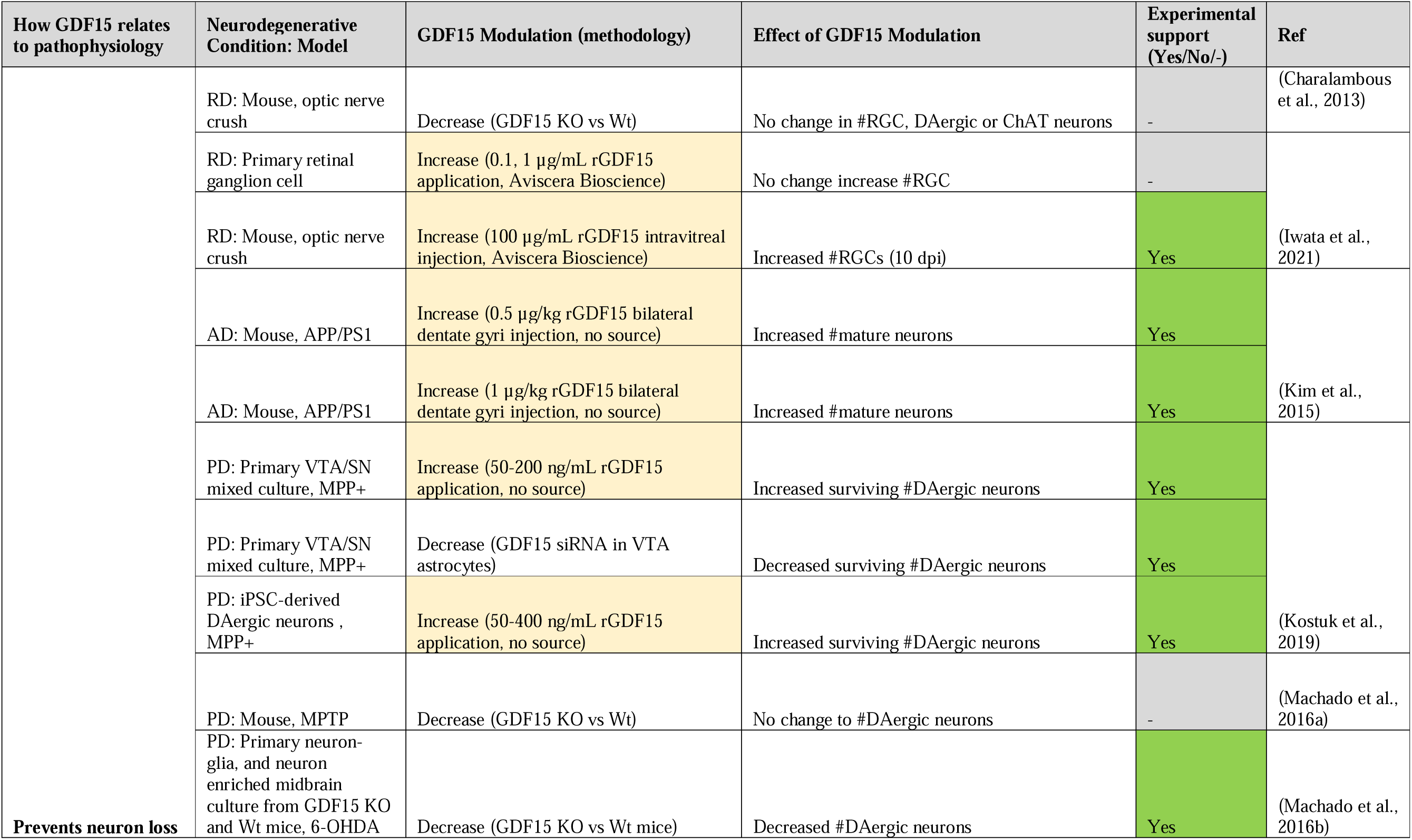
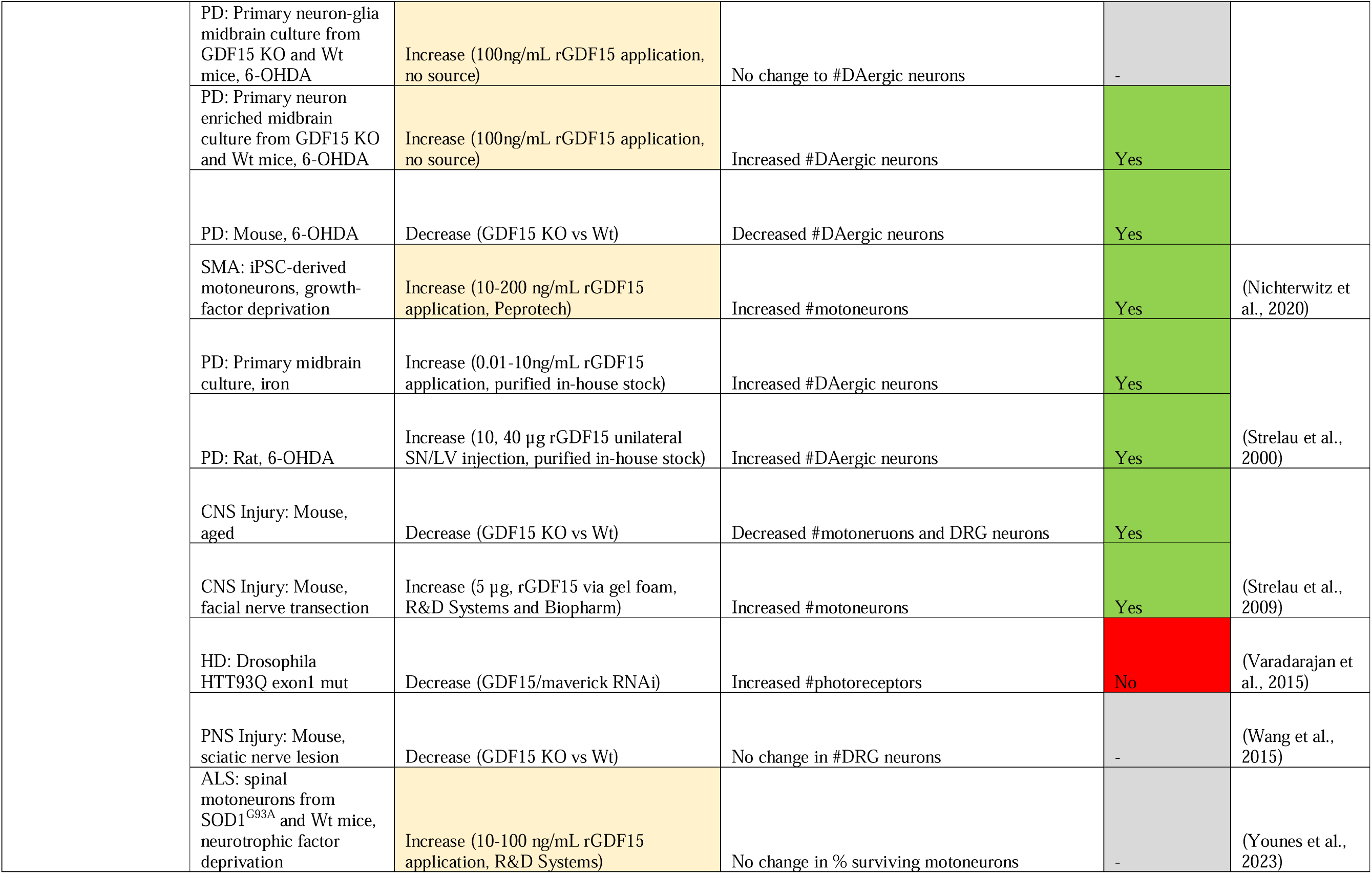
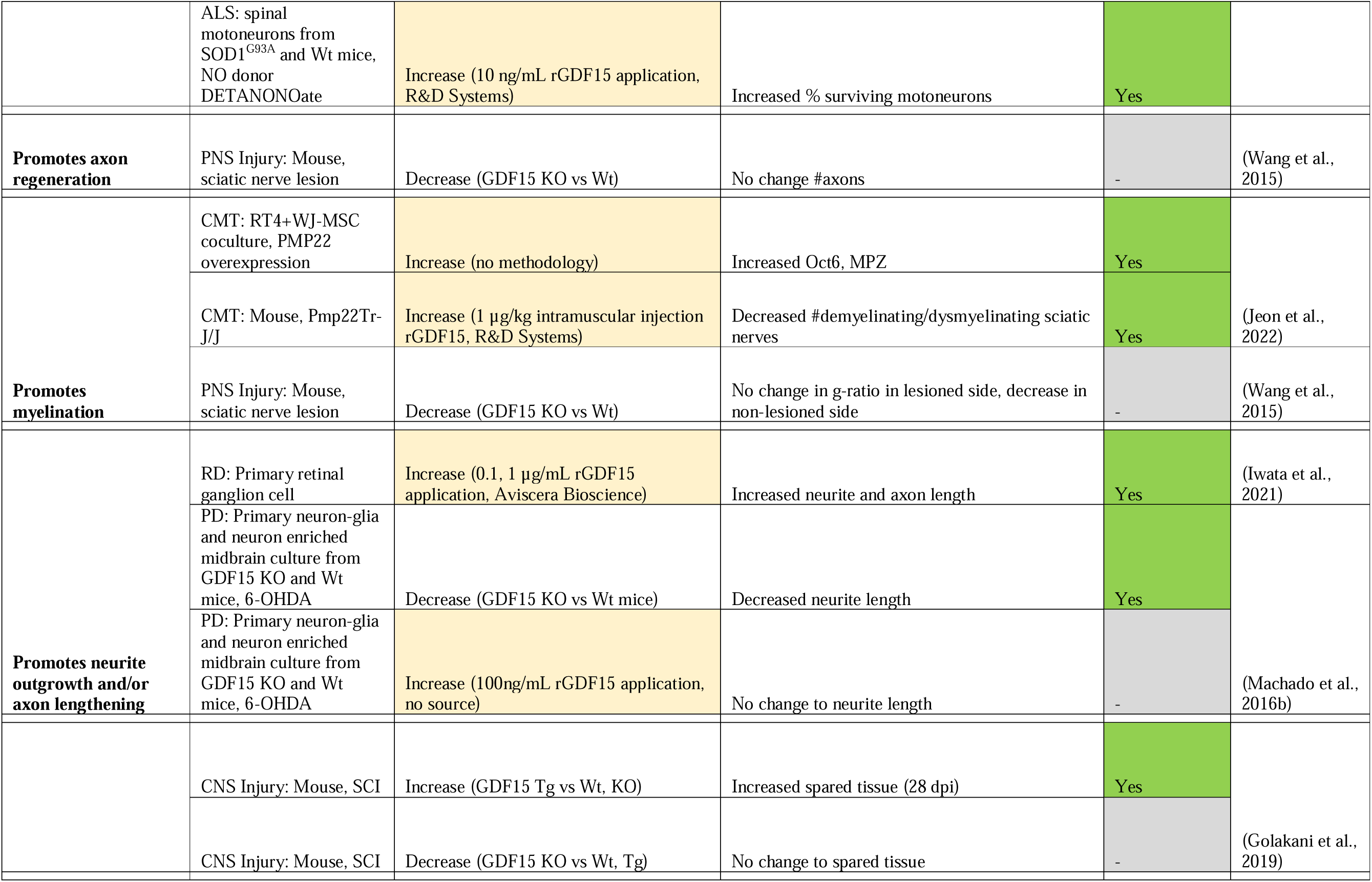
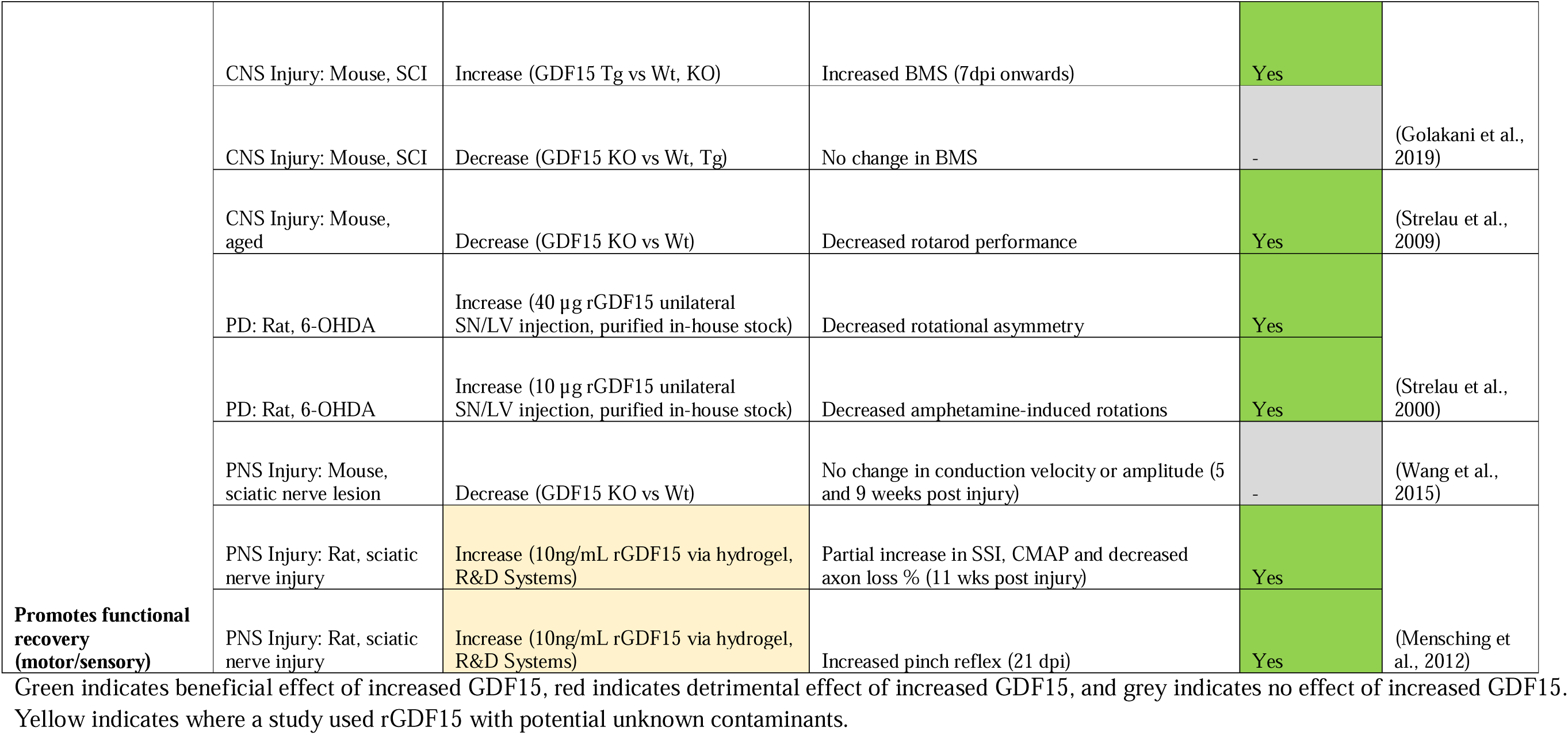
Modulating GDF15 in ND models is neuroprotective by promoting cellular regeneration and functional recovery.

The above studies attributed changes in functional outcomes following induced injury/insult to the extent of neuron loss, however, the evidence linking this to GDF15 is mixed (Table 7). In support of GDF15’s involvement, aged GDF15 null mice have fewer motoneurons and dorsal root ganglia neurons than their wildtype counterparts (Strelau et al., 2009). Another study showed loss of dopaminergic neurons and decreased neurite length following 6-OHDA treatment in GDF15 knockout mice and their midbrain neuron-glia cultures relative to wildtype (Machado et al., 2016b). rGDF15, however, rescued dopaminergic neurons in 6-OHDA treated rats and iron-treated midbrain cultures (Strelau et al., 2000). Furthermore, GDF15 overexpression in SCI mice decreased the area of injured spinal cord tissue at 28 dpi (Golakani et al., 2019). However, the remaining studies knocking down or silencing GDF15 do not consistently replicate these effects. In fact, one study showed silencing GDF15 was protective in a drosophila model of Huntington’s disease, by ameliorating endoplasmic reticulum-stress induced apoptosis in photoreceptors (Varadarajan et al., 2015). In mice, neuron loss did not vary between knockout and wildtype genotypes following sciatic nerve lesion (Wang et al., 2015), optic nerve crush (Charalambous et al., 2013), or MPTP treatment (Machado et al., 2016a). Further, GDF15 knockout had no effects on injury size (Golakani et al., 2019), axon number, or g-ratio as a measure of demyelination (Wang et al., 2015). Interestingly, when grown in a co-culture, GDF15 silencing of VTA astrocytes potentiated dopaminergic neuron loss after MPP+ treatment (Kostuk et al., 2019). This suggests that although astrocytes may contribute to GDF15-mediated neuron preservation in mixed neuron-glia culture, compensatory mechanisms *in vivo* may offset the effects of GDF15 silencing, explaining the lack of exacerbated neuron loss. Further research would benefit from replicating these models using transgenic or pharmacological GDF15 expression to fully uncover its protective effect.

### 3.5. GDF15 mediates peripheral rather than local neuroinflammatory responses

Multiple studies included in this review examined the effect of GDF15 on inflammation and immune responses in neurodegenerative models (Table 8). First and foremost, several studies indicated changes in cytokine expression in response to GDF15 modulation. GDF15 overexpressing mice with SCI showed significantly increased CCL2 expression, but not IL-6 at 28 dpi (Golakani et al., 2019). Contrastingly, while GDF15 knockout showed no change in CCL2, IL-6 and other signature molecules IL-1β, MAC-2, Arg-1 and Ym1 were elevated in the first 0 – 7 days in nerve tissue distal to crush injury (Wang et al., 2015). Another study found 6-OHDA-treated GDF15 knockout mice had increased IL-6 and iNos at 6 and 14 dpi in the caudate putamen and substantia nigra respectively, with no change in TGF-β1 or TNF-α (Machado et al., 2016b). However, limited changes were identified following MPTP injection. Cytokines iNos, TNFα, IL-6, Arg1, Fizz-1, Ym-1 and TGFβ-1 were unchanged between GDF15 null and wildtype mice until 90 dpi, when they were downregulated (Machado et al., 2016a). This milder cytokine response could be linked to the relatively transient nature of MPTP neurotoxicity (Machado et al., 2016a), however, without consistent timepoints between different models, this is speculative. Nevertheless, GDF15’s role in altering cytokine expression following neurodegenerative injury or insult was further indicated by *in vitro* studies. Following 6-OHDA treatment, IL-6 was more elevated in primary mixed cultures derived from GDF15 knockout mice (Machado et al., 2016b). SH-SY5Y neurons treated with Aβ42 expressed less TNFα, IL-6, IL-1B and IL-8 when GDF15 was overexpressed (Xiong et al., 2023). When exposed to siGDF15 treated MSC-derived exosomes, the opposite effect was observed in these neurons (Xiong et al., 2021). Overall, although modulation of GDF15 alters cytokine expression profiles in response to induced neurodegeneration, lack of replicability in the types of cytokines and the timepoints measured between these studies makes an overall pattern difficult to ascertain.

**Table 8.**
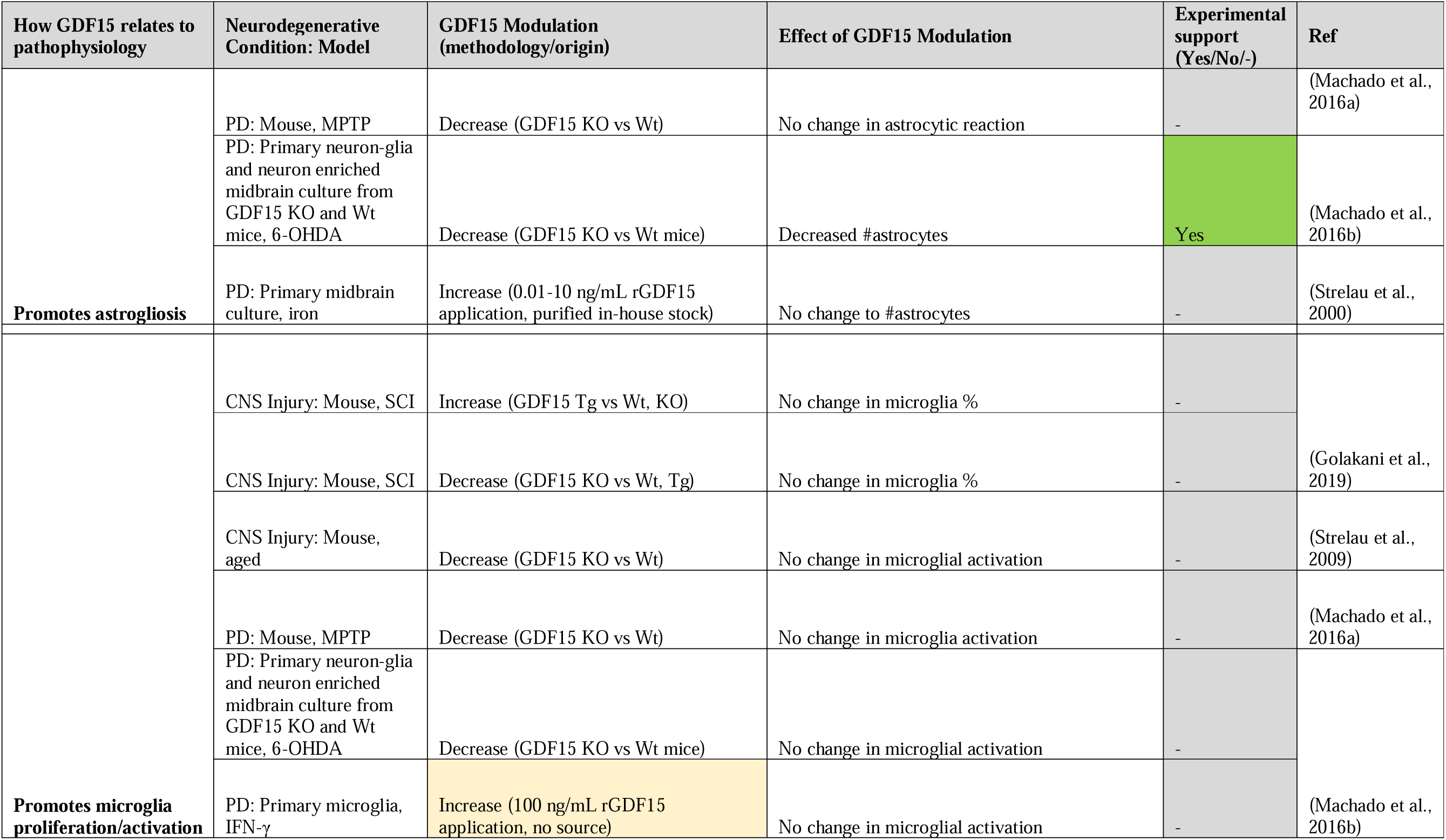
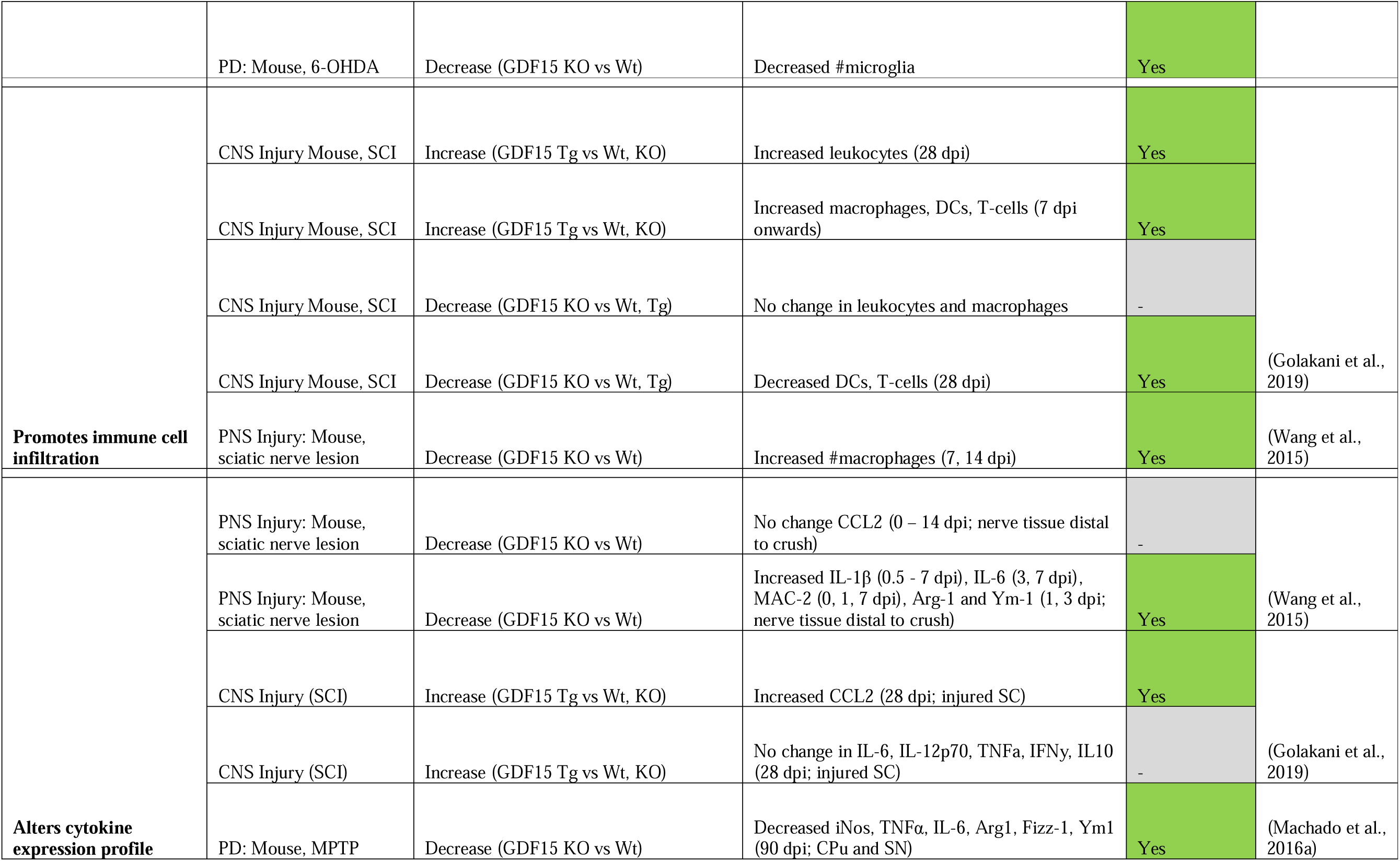
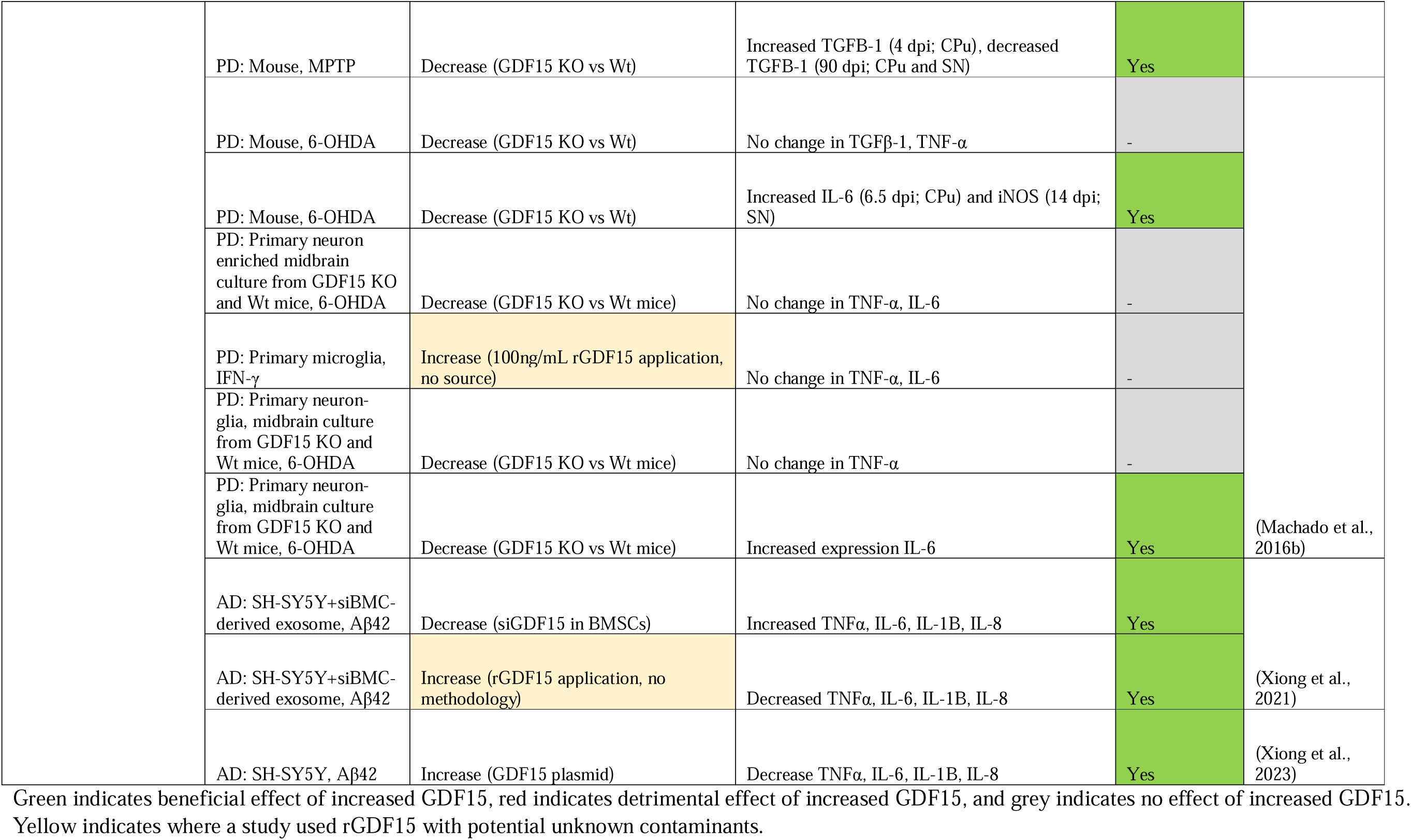
Modulating GDF15 in ND models is neuroprotective by mediating inflammation.

Secondary to cytokine dysregulation, our review identified two *in vivo* studies that examined how GDF15 may influence peripheral immune infiltration in response to induced injury mechanisms (Table 8). One study identified an altered immune cell response in GDF15 transgenic and knockout mice following SCI. Macrophages, dendritic cells, and T-cells were elevated from 7 dpi, and leukocytes from day 28 in the injured spinal cord of GDF15 overexpressing mice. GDF15 null mice however, had reduced dendritic and T-cell infiltration, and unchanged macrophage and leukocyte levels (Golakani et al., 2019). The other study examined responses in the PNS, where GDF15 knockout increased macrophages distal to the injury in the 1 – 2 weeks following sciatic nerve lesion (Wang et al., 2015). As macrophage recruitment was increased in one GDF15 null injury model and not the other, these differences may relate to inherent distinctions between the peripheral and central nervous system, though further experimental work is required to support this assertion.

Neuroinflammation underlying neurodegeneration is associated with glial activity and proliferation. In fact, two included studies showed that astrocytes express and secrete GDF15 in response to induced neuroinflammation (Yi et al., 2015, Lee et al., 2017). However, only one of our included studies demonstrated an effect of modulating GDF15 on these responses (Table 8). 6-OHDA treated-GDF15 knockout mice had reduced numbers of microglia (Machado et al., 2016b). This same study indicated substantial astrocyte loss in mixed midbrain cultures derived from the same mouse line (Machado et al., 2016b). In contrast, other studies did not identify a change in astrocyte reactivity (Machado et al., 2016a), microglial activation (Strelau et al., 2009) or overall numbers of these glial cells (Strelau et al., 2000, Golakani et al., 2019). Therefore, there is limited evidence linking exogenous GDF15 modulation to glial neuroinflammatory responses following induced neurodegeneration.

### 3.6. Elevated GDF15 is associated with neurodegenerative disease and injury

The second aim of this systematic review was to assess GDF15 levels as a biomarker in individuals with neurodegenerative disease or injury. Overall, the studies examined show that GDF15 levels are elevated in some neurodegenerative conditions and are associated with disease severity and incidence (Table 9; Supplementary Table 2).

**Table 9.**
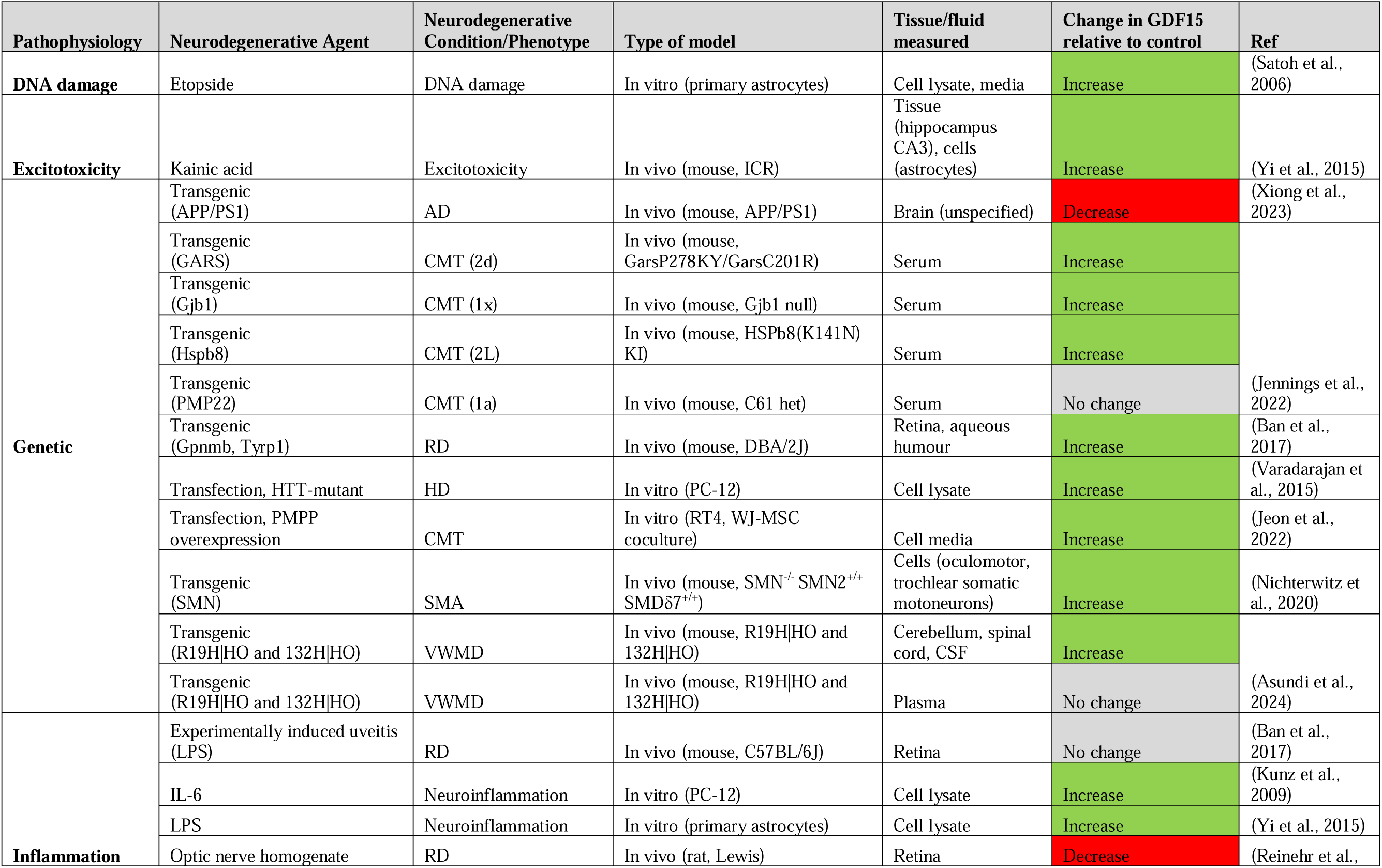
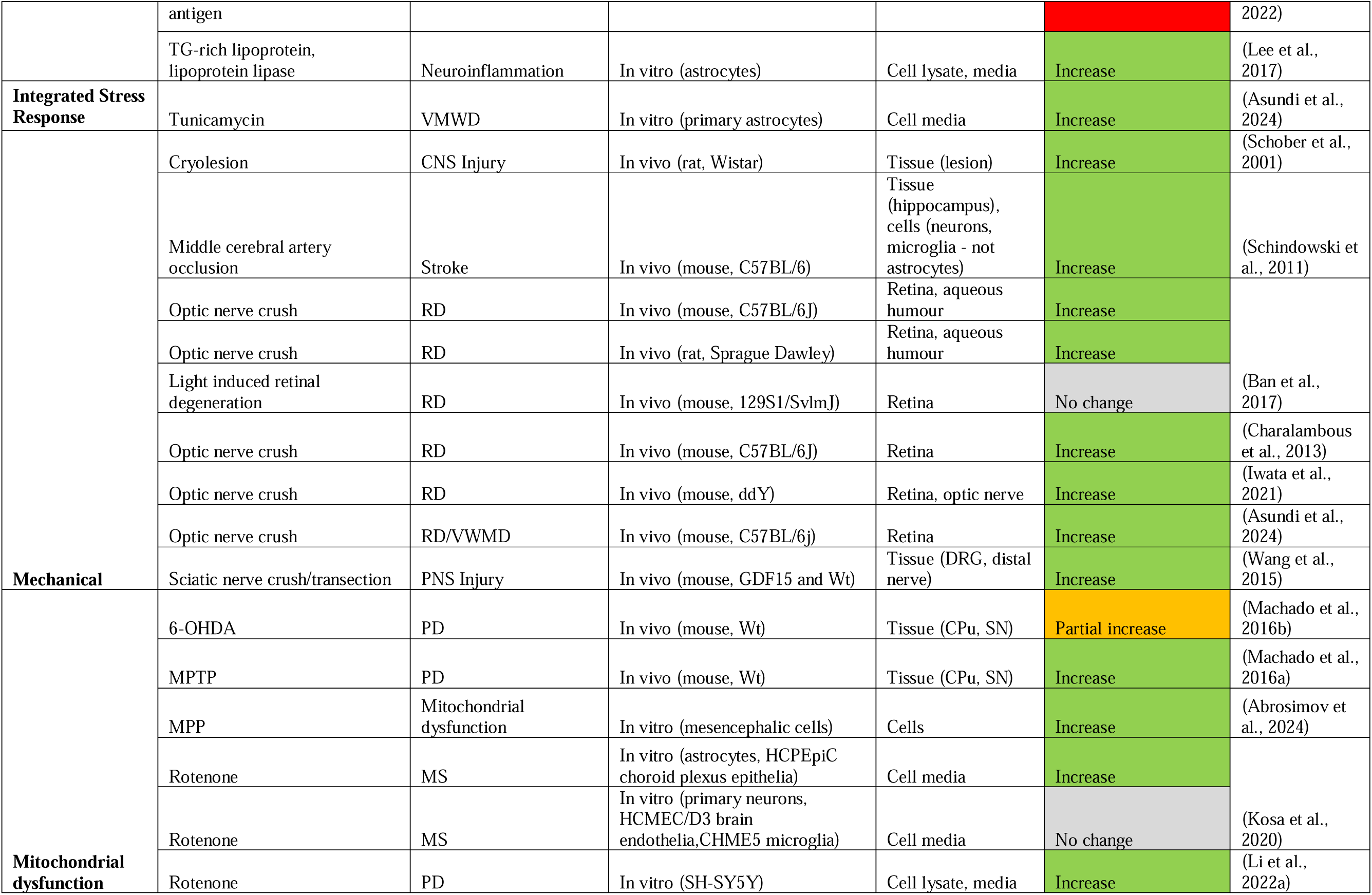
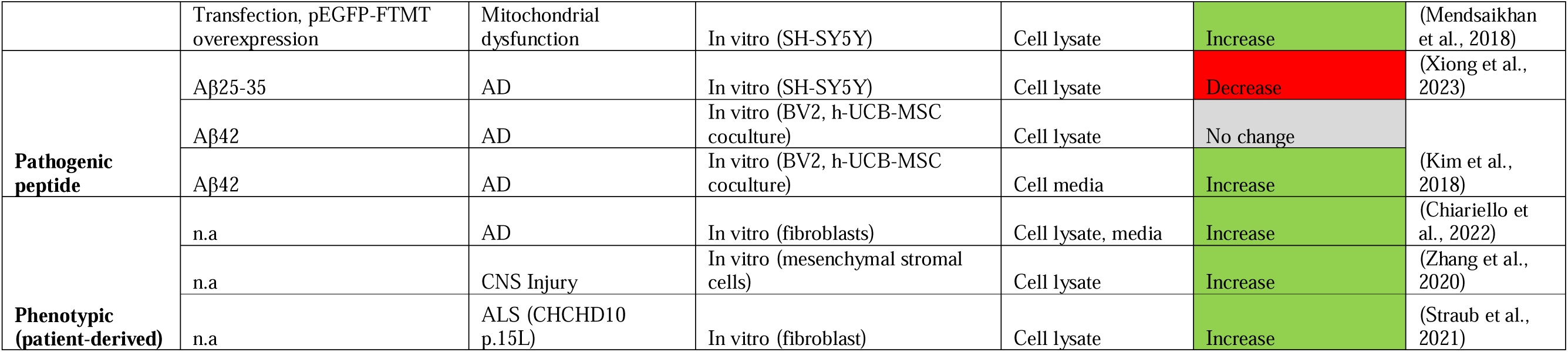
Measuring GDF15 expression levels in models of neurodegeneration.

#### Mitochondrial disease

Mitochondrial diseases are caused by mutations in mitochondrial DNA (mtDNA) and mitochondrial genes (Zhang et al., 2024) and show consistent upregulation of GDF15. As a secreted protein, GDF15 circulates in blood in a normal range of (200 – 1200 pg/mL) (Breit et al., 2021). Across paediatric and adult cases, blood levels of GDF15 were elevated in mitochondrial disease patients compared to their healthy counterparts (Montero et al., 2016, Ji et al., 2017, Maresca et al., 2020, Miyaue et al., 2020b, Miyaue et al., 2020a, Huddar et al., 2021, Peñas et al., 2021, Li et al., 2022b, Riley et al., 2022, Yatsuga et al., 2015, Jean et al., 2022). Two of these studies found that this elevation related to neurodegenerative outcomes. The first highlighted an association between GDF15 levels and supratentorial grey matter atrophy and white matter microstructural changes identified by magnetic resonance imaging (MRI) (Evangelisti et al., 2022). The second showed that this relationship is further strengthened by stratifying cases into mitochondrial encephalomyopathy with lactic acidosis and stroke-like episodes and myoclonic epilepsy with ragged red fibres (Maresca et al., 2020). However, this was not replicated in a broader cohort (Li et al., 2022b), and remains unclear. Although GDF15 is associated with aging, neurodegeneration and all-cause mortality more generally, it should be noted that increased levels were not associated with age in mitochondrial disease (Montero et al., 2016, Nohara et al., 2019, Maresca et al., 2020, Miyaue et al., 2020a, Huddar et al., 2021), with one weak exception in paediatric cases (Li et al., 2022b). This suggests that GDF15 elevation in these conditions is not an age-associated phenomenon, rather, reflective of an upregulated mitochondrial stress response. Indeed, mitochondrial disease patients had higher circulatory GDF15 levels compared to non-mitochondrial neuromuscular disease controls (Montero et al., 2016, Ji et al., 2017, Huddar et al., 2021, Peñas et al., 2021, Li et al., 2022b, Riley et al., 2022). Further, several studies utilised mitochondrial dysfunction to induce a neurodegenerative phenotype *in vitro* and *in vivo*, which caused elevated GDF15 expression and secretion (Machado et al., 2016a, Machado et al., 2016b, Kosa et al., 2020, Li et al., 2022a, Mendsaikhan et al., 2018, Abrosimov et al., 2024). Therefore, heightened circulatory GDF15 indicates mitochondrial stress and dysfunction, likely associated with neurodegenerative pathophysiology.

#### Alzheimer’s disease and related dementias

Upregulated blood GDF15 was further revealed to be associated with risk of dementias, neurodegenerative diseases affecting cognition, including AD. AD is characterised by progressive neuron and synapse loss, alongside accumulation of amyloid-β (Aβ) plaques and hyperphosphorylated Tau neurofibrillary tangles (Chiariello et al., 2022). With respect to relative levels in plasma, GDF15 was increased in AD (Chai et al., 2016) and dementia patients (Ji et al., 2023) compared to individuals with no cognitive impairment. A significant increase was also reported in individuals with mild cognitive impairment (Chai et al., 2016, Jiang et al., 2015, Ji et al., 2023). In contrast with blood GDF15, CSF levels did not vary in AD patients compared to controls (Chiariello et al., 2022). In assessing risk, one study performed 2-sample Mendelian randomization analysis using summary-level datasets from GWAS for an AD population. The authors identified that increased serum GDF15 levels was associated with higher AD risk (Wu et al., 2021). This relationship was further evident in assessing prospective cohorts for AD and dementia incidence (McGrath et al., 2020, Walker et al., 2023, You et al., 2023, Lu et al., 2022, Guo et al., 2024), although significance was lost for AD when including N-terminal pro-brain natriuretic peptide (NT-proBNP), a marker of ventricular distention (McGrath et al., 2020). Another study found a correlation between elevated GDF15 and AD Pattern Similarity (AD-PS) score, an index of neuroanatomical dementia risk that compares grey matter spatial patterns (Casanova et al., 2023). Consistent with this, other examinations showed increased peripheral GDF15 was associated with white matter hyperintensities (Chai et al., 2016) and that this was strengthened in cohorts with cognitive impairment relative to cognitively normal individuals (Jiang et al., 2015).

In addition to its potential as an AD blood biomarker, dysregulation of central GDF15 was reported, albeit with inconsistent findings. Firstly, increased GDF15 was identified in multiple post-mortem AD brain areas (Wang et al., 2022). This finding conflicted with another study that reported no change in GDF15 levels in post-mortem frontal cortex tissue (Chiariello et al., 2022). Although, this study did find an increased ratio of mature to precursor GDF15 in AD patients and centenarians. Given that processed, mature GDF15 rapidly diffuses away into circulation (Bauskin et al., 2005), this change may reflect an age or disease-associated increase in GDF15 processing. However, dermal fibroblasts taken from AD subjects expressed significantly more GDF15 mRNA and secreted more protein into media than age-matched non-demented subjects (Chiariello et al., 2022). This incongruity implies that increased GDF15 is more closely associated with Alzheimer’s pathology than with healthy aging. Additionally, BV2 microglia secreted more GDF15 into media with MSCs when incubated with Aβ42 (Kim et al., 2018), suggesting GDF15 upregulation is triggered by AD pathology to attenuate pathogenic mechanisms. On the other hand, one study found that GDF15 mRNA and protein was reduced in both the brain of a familial AD mouse model compared to wildtype controls, and in SH-SY5Y cells incubated with Aβ 25-35 peptide (Xiong et al., 2023). These inconsistencies in GDF15 expression may relate to differences in microglia and neuronal responses to Aβ, as well as variance between familial and late onset AD pathology, although these effects are not compared experimentally. Ultimately, whilst GDF15 upregulation in AD pathogenic models and *ex vivo* tissue is inconsistently demonstrated, it may have potential utility as a biomarker for AD and other dementias.

#### Synucleinopathy

Synucleinopathies represent a class of neurodegenerative diseases pathologically characterised by aggregated and phosphorylated α-synuclein. These diseases include Parkinson’s disease (PD), PD with dementia (PDD), Dementia with Lewy Bodies (DLB) and Multiple System Atrophy (MSA) and vary clinically by variation in region and cellular pathology. Two studies identified that serum GDF15 was elevated in PD cases compared to controls (Yao et al., 2017, Miyaue et al., 2020b), whilst another showed no statistical difference (Davis et al., 2020). One study examined MSA patients (Yue et al., 2020), demonstrating elevated serum GDF15 relative to both healthy controls and PD. Further, GDF15 levels were higher in cases with longer disease duration, older age, and MSA-Parkinsonian subtype. A sex effect was also uncovered, with higher serum GDF15 in male synucleinopathy patients relative to females (Yao et al., 2017, Yue et al., 2020). With respect to CSF measurements, GDF15 was increased in all Lewy body diseases, particularly for the PDD subgroup (Maetzler et al., 2016). Older age at symptom onset was related to higher GDF15 in two studies (Maetzler et al., 2016, Davis et al., 2020) but not in another (Yao et al., 2017). Furthermore, elevated serum GDF15 was associated with disease duration and worsened motor and cognitive scores and was reflective of poorer neurological outcomes in these synucleinopathy cohorts (Maetzler et al., 2016, Yao et al., 2017). Further, measures of neuronal death and axonal damage t- and p-Tau correlate with GDF15 levels in CSF (Maetzler et al., 2016). A recent longitudinal assessment of a large preclinical group showed plasma GDF15 levels as associated with higher risk for PD (You et al., 2023), whilst an earlier study did not identify this risk (Wu et al., 2021). Induction of parkinsonian-like pathology in cell and animal models was also found to trigger upregulated GDF15. Use of inflammatory neurotoxins increased GDF15 expression in the caudate putamen and substantia nigra of mice (Machado et al., 2016a, Machado et al., 2016b), and in both cell lysate and media of SH-SY5Y neurons (Li et al., 2022a). Thus, there is some support for increased serum GDF15 in synucleinopathy, and this elevation may be an intrinsic component of synuclein pathogenesis.

#### Multiple Sclerosis and Neuromyelitis Optica Spectrum disorder

MS is a chronic neuroinflammatory disorder causing demyelination and neuron degeneration in the entire CNS. MS is further categorised into clinical phenotypes that include relapsing-remitting (RRMS), characterised by acute and highly variable attacks; primary progressive (PPMS), which gradually worsens from onset; and secondary progressive (SPMS), which initially presents as RRMS and progresses to resemble PPMS later in disease development (Kuhlmann et al., 2023). Studies that do not differentiate between MS cases fail to detect any differences in serum GDF15 relative to healthy controls (Yatsuga et al., 2015, Melvin et al., 2019). Interestingly, this is not the case when studies examine the specific clinical phenotypes of MS. Individuals diagnosed with PPMS or SPMS had elevated GDF15 in serum and CSF relative to control (Kosa et al., 2020, Baeva et al., 2023) and RRMS cases (Kosa et al., 2020). No significant increase was observed between RRMS and controls in these studies (Kosa et al., 2020, Baeva et al., 2023). Given the heterogeneity in MS cases, it is therefore unsurprising that altered levels of GDF15 was found to relate to clinical course and severity. In cohorts with undefined MS, serum GDF15 was elevated 7.4 months into disease progression (Melvin et al., 2019), and positively associated with Expanded Disability Status Scale (Nohara et al., 2019). However, this could not be replicated in a purely PPMS group (Baeva et al., 2023). In fact, baseline GDF15 was reduced in PPMS patients who had worsened motor outcomes at 18 months (Baeva et al., 2023). Contrastingly, GDF15 levels were higher in RRMS patients who did not develop gadolinium-enhancing or T2 lesion activity, indicative of focal inflammation (Kuhlmann et al., 2023, Amstad et al., 2020). When put together, these results suggest that the overall pattern of GDF15 dysregulation in MS is possibly linked to both clinical and pathological phenotypes and may be reflective of underlying neuroinflammation. Future studies examining GDF15 as an MS biomarker should substantiate these findings by directly relating the disease subtype.

Further to fluid biomarker examination of GDF15 levels in MS, there is some support for the role of astrocytes in the production of this cytokine to mediate neuroinflammatory responses. One study found 1 – 10 % of reactive astrocytes and macrophages/microglia expressed GDF15 immunoreactivity in demyelinating lesions localised to the frontal cortex of a patient with SPMS (Satoh et al., 2006). Cell models recapitulating MS pathology have shown that astrocytes express and secrete greater GDF15 in response to oxidative stress (Satoh et al., 2006) and mitochondrial dysfunction (Kosa et al., 2020). This response is likely to both induce neurotrophic effects in damaged neurons and signal cell stress associated with MS lesions.

Research examining serum GDF15 in individuals with Neuromyelitis Optica Spectrum disorder, a demyelinating autoimmune condition akin to MS, was also included in this review. No significant differences were shown between patient and control cases (Miyaue et al., 2020a, Yatsuga et al., 2015), suggesting it is not a suitable biomarker for this disease and potentially reflecting variation from MS pathophysiology.

#### Glaucoma

Glaucoma refers to a group of neuroretina degenerative diseases associated with retinal ganglion cell death. Studies included in this review assessed GDF15 in the context of biomarkers for primary open angle glaucoma and pseudoexfoliative glaucoma, showing elevated levels in both diseases (Ban et al., 2017, Lin et al., 2020). Further, this elevation in GDF15 appears to be unrelated to surgical interventions, with one study showing some patients have decreased and others increased GDF15 post-operatively (Lin et al., 2021). In rodent models, GDF15 expression changes were largely dependent on the type of model used. GDF15 was upregulated both in the retina and aqueous humour of chronic glaucoma mouse models at 1 year (Ban et al., 2017). Further, optic nerve crush also elevated GDF15 levels in mice (Charalambous et al., 2013, Iwata et al., 2021, Asundi et al., 2024) and rats (Ban et al., 2017) in 1 – 9 dpi. In contrast, no change was observed at any timepoint in the mouse retina following light-induced retinal degeneration or in experimentally induced uveitis (Ban et al., 2017). Moreover, 28 days after systemic injection of optic nerve homogenate antigen, mRNA expression of this cytokine was diminished in the inner retinal layer (Reinehr et al., 2022). Therefore, while GDF15’s association with retinal degeneration varies, fully understanding its role in neuroretina inflammation requires further evidence and additional timepoints.

#### Motor neuron disease

Research examining GDF15 dysregulation in MNDs is limited, with conflicting support between human and animal/cell-based studies. With respect to ALS, serum GDF15 was not associated with disease risk (Wu et al., 2021). However, fibroblasts derived from an ALS patient with a CHCHD10 p.15L gene variant were shown to express more GDF15 than the wildtype line (Straub et al., 2021). Similarly, one study showed no significant serum difference between Spinal Muscular Atrophy Jokela type individuals and healthy controls (Järvilehto et al., 2022). On the other hand, in a mouse model of this condition, examination of motoneurons that are resilient to degeneration showed differential upregulation of GDF15 alongside other anti-apoptotic factors (Nichterwitz et al., 2020). Thus, although there is insufficient evidence to assert if it acts as a suitable biomarker for either MND, GDF15 may be associated with their underlying neuropathogenic processes either by reflecting cell stress or promoting cell survival pathways.

#### Charcot-Marie-Tooth disease (CMT)

CMT is a type of demyelinating, peripheral nervous system neuropathy. One study determined serum GDF15 was elevated across all subtypes of this condition and was worsened by disease severity as measured by the progression score CMTES (Jennings et al., 2022). A recent study indicated GDF15 did not vary between genetic variants CMT1a and PMP22 for the disease (Palu et al., 2023). This is supportive of findings from mouse models of CMT that show GDF15 was elevated in serum (Jennings et al., 2022) and in schwannoma cell media when transfected with PMP22 (Jeon et al., 2022).

#### Huntington’s disease (HD)

HD is a complex neurodegenerative disorder with psychological, cognitive, and motor symptoms. Pathologically, this disease is driven by mutant HTT fibrillation and multimerization. Our review did not identify any HD biomarker studies, although the HTT mutant was shown to lead to higher levels of GDF15 than control transfected neurons (Varadarajan et al., 2015).

#### Vanishing White Matter Disease (VMWD)

VWMD is a progressive hypomyelinating disease caused by bi-allelic variants in eukaryotic initiation factor 2B, which mounts the integrated stress response. One included publication examined VWMD in human populations, with complementary *in vivo* and *in vitro* modelling (Asundi et al., 2024). Elevated GDF15 was reported in VWMD patient CSF compared to controls, but not in plasma or serum. This was consistent with VWMD mouse models, which additionally exhibit increased GDF15 expression in the cerebellum and spinal cord at 4 months relative to wildtype. Upregulated GDF15 was determined to be astrocytic in origin; GDF15 transcripts were localised to affected mouse forebrain astrocytes, and increased secretion occurred in response to tunicamycin *in vitro* (Asundi et al., 2024). In summary, this study reveals a significant association between VMDW and elevated GDF15, mediated by the integrated stress response.

#### Neurodegeneration secondary to injuries to the central and peripheral nervous system

Studies examining GDF15 in the context of secondary neurodegeneration due to neurotrauma were also assessed in this review. One key finding was that circulatory GDF15 is closely related to the incidence, severity, and risk of stroke. Increased GDF15 in blood was observed in the acute phase following an ischemic event in affected individuals relative to healthy controls (Lu et al., 2020, Liu, 2021, De Marchis et al., 2023). It was also evident that blood GDF15 at time of admission was consistently elevated in stroke patients who later displayed poorer modified Rankin Scores (mRS) both at discharge and 90 days follow up (Worthmann et al., 2011, Jeong et al., 2020). Elevated blood GDF15 was furthermore associated with injury/inflammation biomarkers glial protein S100 calcium binding protein B and IL-6 (Lu et al., 2020, Worthmann et al., 2011) and with lesion size and severity (Liu, 2021, Lu et al., 2020, Jeong et al., 2020). Interestingly, GDF15 on admission correlated with depression scores at 90 days and was predictive of post-stroke associated depression (Lu et al., 2020). Longitudinal studies linked blood GDF15 levels with the risk of incident stroke/TIA, although there was some evidence this was related to cardiovascular events (You et al., 2023, De Marchis et al., 2023, Andersson et al., 2015, Lind et al., 2015).

There is some evidential support for central as well as peripheral expression of GDF15 in response to ischemic stroke. Increased GDF15 staining was identified in a small subset of reactive astrocytes and macrophages/microglia in post-mortem ischemic lesions following acute cerebral infarction (Satoh et al., 2006). Furthermore, GDF15 expression and immunoreactivity was also elevated in mouse hippocampus 3 – 24 h following middle cerebral artery occlusion (Schindowski et al., 2011). Together with blood biomarker findings highlighted above, these studies indicate that GDF15 serves as a putative stroke biomarker, although future work should elaborate further on the source of centrally induced GDF15.

Besides stroke, examination of GDF15 in neurotrauma-induced degenerative conditions in human populations was limited. One study showed that serum GDF15 was elevated in patients following subarachnoid haemorrhage, and this elevation in the first 9 days related to worsened primary and secondary neurological outcomes at 90 day follow up (Csecsei et al., 2023). Another study examined mesenchymal stromal cells derived from individuals with SCI or traumatic brain injury (TBI). The authors identified increased GDF15 secretion into cell lysate, indicative of a widespread cell stress response (Zhang et al., 2020). This induction was also reported in injured rodent CNS and PNS tissue in response to cryolesion (Schober et al., 2001) and sciatic nerve crush (Wang et al., 2015) respectively. Overall, although limited in number and scope, these studies highlight that increased expression of GDF15 can be identified in mixed neurotrauma states.

## 4. Discussion

To better understand GDF15’s role in neurodegeneration, we conducted a systematic review of the literature. Our review explored the possible mechanisms underlying GDF15’s neuroprotective function in animal and cell models, as well as its potential as a biomarker for neurodegenerative conditions. Overall, the included studies showed that exogenously increasing GDF15 is generally neuroprotective, whilst silencing/knockdown is linked to largely exacerbated neurodegeneration. Favourable outcomes related to GDF15 modulation were attributable to its anti-apoptotic effects, increased cell viability, alleviation of mitochondrial and oxidative stress, and modulation of neuroinflammation. Furthermore, biomarker studies show that GDF15 levels were higher in individuals with most chronic neurodegenerative conditions, and, with the exception of MS, these elevated levels were typically associated with increased disease incidence and severity. Together, this highlights the potential utility of GDF15 as both a therapeutic tool and a possible biomarker in neurodegeneration.

### 4.1. The U-curve dose-response relationship of GDF15 in neurodegeneration

Our review highlights that GDF15 acts as a neuroprotective agent in disease and injury models, yet its elevation in some patients is linked to poorer outcomes. This is consistent with existing literature on GDF15’s role in exercise and cancer, demonstrating a U-shaped dose-response to stress. Vigorous endurance exercise, which induces transient mitochondrial stress, spikes plasma GDF15 comparable to patients with MD, infection, and cancer (Klein et al., 2021, Conte et al., 2020). This increase in GDF15 is also observed in mice; however, unlike the effect of high-dose exogenous delivery of rhGDF15, it does not suppress appetite and running activity (Klein et al., 2021). Higher endogenous GDF15 is also observed in chronically inactive and aging populations (Conte et al., 2020) and is inversely associated with survival (Conte et al., 2018). Furthermore, while GDF15 offers protection by alleviating impaired mitochondrial function, its induction of the mitochondrial integrated stress response in papillary thyroid carcinoma patients can also contribute to cancer progression via the GDF15-STAT3 pathway (Kang et al., 2020). Therefore, optimal levels of GDF15 exhibit protective and therapeutic effects, but elevated or sustained levels in specific pathogenic contexts indicate insurmountable cellular stress and serve as disease biomarkers (Figure 2). Crucially, elevated GDF15 observed in disease models and in epidemiological studies does not imply disease causality. As proposed by Breit and colleagues, it is possible that while neurodegeneration induces GDF15 in proportion to disease severity as a reparative response, the prolonged elevation of GDF15 eventually becomes inadequate in alleviating or reversing disease-related stress (Tsai et al., 2018, Breit et al., 2021). This inverted u-curve relationship may explain why chronically induced endogenous GDF15 in affected individuals fails to lessen aspects of neurodegeneration that are more effectively addressed in cell and animal models. Supporting this, one included study showed that administering 2Bact, an inhibitor of the integrated stress response, to Vanishing White Matter disease and optic nerve crush mice resulted in reduced GDF15 levels and improved pathology (Asundi et al., 2024). Future investigations should prioritise a qualitative analysis of optimal GDF15 levels in neurodegenerative settings.

**Figure 2.**
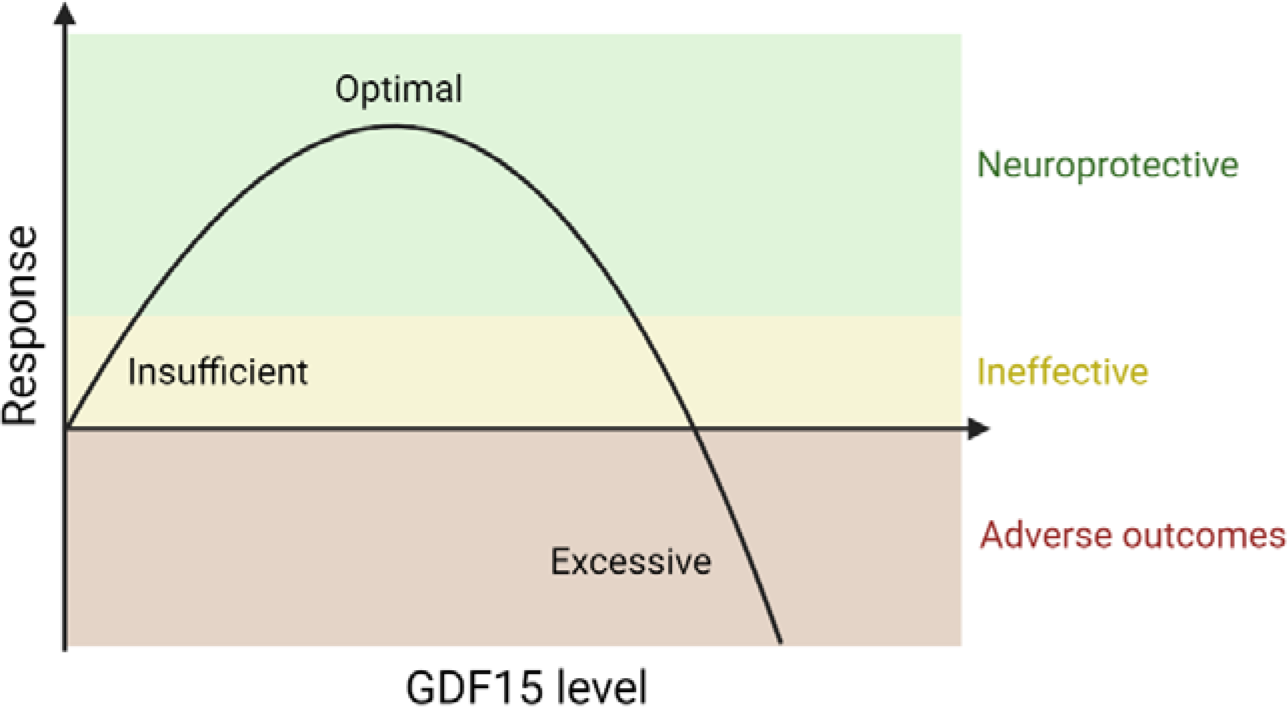
Proposed Inverted U-Curve Dose-Response Relationship of GDF15 to Neurodegeneration. This schematic illustrates the dose-response relationship between GDF15 levels and neurodegenerative outcomes. Low level, basal GDF15 has minimal impact on neurodegeneration. An optimal, intermediate level of GDF15 promotes therapeutic effects and neuroprotection. High levels of GDf15 indicate excessive cell stress and is associated with adverse neurodegenerative effects.

### 4.2. GDF15’s neuroprotective role is likely mediated centrally by GFRAL-RET

Despite the emphasis on the activation of the PI3K/Akt/ERK pathway as a potential downstream pathway of GDF15, it remains unclear how GDF15 triggers this activation in neurodegeneration. Prior to identifying GFRAL as its target receptor, the literature primarily attributed these responses to TGF-β family receptors I (TGF-βI) and II mediated Smad pathway phosphorylation. However, there is no strong evidence to show that these receptors are activated by GDF15 (Yang et al., 2017), as studies show this could be inadvertently reporting TGF-β activity due to contamination of commercial GDF15 preparations (Olsen et al., 2017). Our review highlights a lack of studies investigating the mechanisms of GFRAL- mediated neuroprotection, possibly due to the limited areas of expression identified for this receptor. It is possible that GFRAL expression is more widespread than currently established, or is inducible by disease, but is not detectable by standard laboratory methods (Breit et al., 2021). Alternatively, as the main alternate transcript of GFRAL lacks a transmembrane sequence (Li et al., 2005), it could serve as a soluble receptor (Breit et al., 2021). It is possible that GDF15 binds this soluble form, enabling RET engagement in cells that do not typically express GFRAL (Tsai et al., 2018). This mechanism bears similarity to that of IL-6 trans-signalling, in which soluble IL-6R can bind to co-receptor gp130, making virtually all cells responsive to IL-6 (Mackiewicz et al., 1992). It is well established that RET tyrosine phosphorylation activates MAPK/ERK, PI3K/Akt and Rac/cJun JNK pathways that are anti-apoptotic (Li et al., 2005) and drive cellular proliferation, differentiation, functioning and more (reviewed in Mahato and Sidorova, 2020, Rochette et al., 2020). One study in our review showed that a RET inhibitor suppressed Akt activation and therefore diminished neurotrophic outcomes attributed to rGDF15 supplementation *in vitro* (Iwata et al., 2021). Trans-signalling could account for the discrepancy between the widespread induction of GDF15 and limited GFRAL expression, however, other receptors or modes of activation cannot be ruled out. For instance, GFRAL knockout mice have similar body weight, fat, and mass to wildtype mice, highlighting the possible presence of non-GFRAL mediated GDF15 activity (Klein et al., 2021), though germline GFRAL knockout may cause other compensatory effects. Alternatively, GDF15 may competitively inhibit activation of receptors that mediate harmful effects. For example, GDF15 hinders recruitment of polymorphonuclear leukocytes into myocardial infarcts and may limit occurrence of cardiac rupture (Kempf et al., 2011). Lastly, we cannot rule out the possibility of indirect induction of TGF-β signalling through GDF15-microRNA interaction (Bloch et al., 2015). However, future studies on GDF15 should prioritise mechanistic investigations of GFRAL-RET signalling in neurodegeneration.

### 4.3. GDF15 deficiency may activate compensatory immunomodulatory effects that drive cellular regeneration in neurodegeneration

Our review revealed a disparity in the immunomodulatory effects observed when GDF15 levels were experimentally raised or lowered in neurodegenerative models, with implications for cellular regeneration and functional recovery. GDF15 overexpressing SCI mice had increased infiltration of macrophages and dendritic cells to SC, as well as improved pathology resolution and motor outcomes (Golakani et al., 2019). This suggests that macrophages may aid in myelin debris clearance (Golakani et al., 2019), while dendritic cells recruit regulatory T-cells that alleviate pathology (Li et al., 2008). This favourable effect is reflected in human MS biomarker cohorts, where elevated peripheral GDF15 is associated with improved neuropathological (Amstad et al., 2020) and motor outcomes (Baeva et al., 2023). Unlike in other neurodegenerative conditions where elevated GDF15 correlates with greater disease severity, this may indicate that certain MS individuals reflect a transient stress response, and GDF15 may effectively resolve these effects in these cases. In contrast with increased GDF15, silencing or knocking down GDF15 did not consistently potentiate tissue damage or functional deficits associated with induced injury or disease. GDF15 null mice did not have worsened tissue damage (Golakani et al., 2019), nerve damage or functional output (Golakani et al., 2019, Wang et al., 2015). This asymmetry could stem from the compensatory increase in IL-6 levels observed when GDF15 is depleted (Machado et al., 2016b, Wang et al., 2015, Xiong et al., 2021, Xiong et al., 2023). IL-6 regulates anti-inflammatory as well as pro-inflammatory responses, as evidenced by its ability to alleviate pathogenic changes in vascular injury (Bonaterra et al., 2012) and in myelin antigen triggered neuroinflammation (Casella et al., 2016). Elevated IL-6 levels in GDF15 null mice following peripheral nerve injury may account for why worsened axon loss and demyelination was not reported here (Wang et al., 2015). Contrastingly, increased GDF15 diminishes IL-6 and alleviates Aβ induced impairments by inhibiting neuronal apoptosis *in vitro* (Xiong et al., 2023). Therefore, IL-6 signalling may compensate for GDF15 deficiency, while elevated GDF15 may minimise pro-inflammatory IL-6 responses; however, due to limited and inconsistent studies on this pattern, our conclusions remain speculative and other beneficial immune mediated responses may be involved.

### 4.4. Limitations of this review and recommendations for future work

It is important to highlight that the broad scope of this review may limit the conclusions drawn. Firstly, we did not incorporate specific neurodegenerative disorders like Alzheimer’s or Parkinson’s disease into our database search. Consequently, some relevant studies such as (Fuchs et al., 2013) may have been overlooked, although this is likely only a sporadic occurrence given our search strategy (including search terms GDF15 and its synonyms) was inclusive of titles and abstracts. Nonetheless, our aim was to evaluate neurodegeneration in a general context, rather than specifically investigating the role of GDF15 within distinct disease or trauma states. Secondly, the literature spanned in this review included highly variable animal and cell studies. This variability extends to factors like method of induced neurodegenerative disease or injury, method of GDF15 modulation, types of outcomes measured, spatial and temporal assessment and animal age, all impacting functional outcomes. For instance, one study shows significant motoneuron loss in GDF15 null mice at 6 months but not at P14 or 3 months (Strelau et al., 2009). This suggests many facets of disease or injury-related responses may not be accounted for in our assessment. Despite this, the consistency in findings across these studies underscores the robustness of GDF15 as a neuroprotective factor. Lastly, risks associated with pharmacological administration of GDF15 were not extensively considered and may impact translation of GDF15 therapeutics into clinical settings. In depth investigations should be undertaken to identify how to best balance the benefits of GDF15 supplementation against its risks, including anorexia/cachexia and weight loss (Breit et al., 2021, Tsai et al., 2018). Further, doses administered in preclinical studies raise GDF15 levels well beyond normal physiological conditions (Klein et al., 2021), questioning their relevance. Future research should consider human-equivalent dosing for more translatable findings.

Results drawn from relevant biomarker studies also have important considerations. Firstly, the included studies may underreport GDF15 levels in individuals with homozygous and heterozygous H6D variants (Karusheva et al., 2022). Unfortunately, our review lacks a straightforward solution to this issue, as it necessitates a direct performance comparison of all available immunoassays. As a result, there is a possibility that many of these studies report inaccurate intergroup differences. This highlights a critical need for biomarker studies to either utilise assays that specifically accommodate for this H6D variant or incorporate suitable statistical models to address variability. Furthermore, the use of prospective cohorts in some studies posed challenges in distinguishing GDF15-related disease risk directly associated with neurodegeneration as opposed to risks associated with all-cause mortality (Moon et al., 2020), infection (Beydoun et al., 2023) or cardiovascular disease (Wollert et al., 2017). Notably, certain studies lost significance when adjusting for cardiovascular risk factors (McGrath et al., 2020, Andersson et al., 2015), indicating that cardiovascular outcomes may play a significant role in driving this relationship. Therefore, considering the widespread induction of GDF15 across various biological systems and in aging, it is improbable for it to serve as a suitable biomarker with single level performance. GDF15’s utility is improved when used in combination with clinical data or additional blood biomarkers (Guo et al., 2024). Although, repeated measures within an individual may still be useful to assess treatment activity wherever available (Brown et al., 2012, Brown et al., 2003, Danta et al., 2017).

## 5. Conclusion

Neurodegeneration represents a complex phenomenon characterised by limited treatment options to impede or decelerate its progression. One of the primary pathophysiological mechanisms implicated in these disorders is neuroinflammation, prompting growing clinical interest in immunomodulatory agents. GDF15 is an anti-inflammatory cytokine with established neurotrophic properties. The evidence outlined in this review suggests that GDF15 expression is induced in response to neurodegenerative pathophysiology and inducing higher levels yields favourable outcomes. Nevertheless, further preclinical investigations which address existing methodological pitfalls are imperative to support these findings mechanistically. Addressing these gaps will facilitate an enhanced understanding of the neuroprotective mechanisms mediated by GDF15, thereby paving the way for novel therapeutic interventions.

## Declaration of Competing Interest

D.A.B. and S.N.B. are inventors on patents owned by St. Vincent’s Hospital that pertain to the clinical use of GDF15. F.I.I, J.F.C, S.T, J.S, V.W.W.T and C.A.F are not aware of any affiliations, memberships, funding, or financial holdings that might be perceived as affecting the objectivity of this review.

## Acknowledgements

Figures were made using Biorender.com and Draw.io.com.

## Funding

This work was supported in part by grants from the National Health and Medical Research Council (Grant #2001350) of Australia, and Medical Research Future Fund (Grant #2017572). C.A.F. is supported by philanthropic funding from the Leece Family Foundation and the Neil & Norma Hill Foundation.

## References

Abrosimov, R., Baeken, M. W., Hauf, S., Wittig, I., Hajieva, P., Perrone, C. E. & Moosmann, B. 2024. Mitochondrial complex I inhibition triggers NAD(+)-independent glucose oxidation via successive NADPH formation, “futile” fatty acid cycling, and FADH(2) oxidation. Geroscience.

Amstad, A., Coray, M., Frick, C., Barro, C., Oechtering, J., Amann, M., Wischhusen, J., Kappos, L., Naegelin, Y., Kuhle, J. & Mehling, M. 2020. Growth differentiation factor 15 is increased in stable MS. Neurol Neuroimmunol Neuroinflamm, 7.

Andersson, C., Preis, S. R., Beiser, A., Decarli, C., Wollert, K. C., Wang, T. J., Januzzi, J. L., JR., Vasan, R. S. & Seshadri, S. 2015. Associations of Circulating Growth Differentiation Factor-15 and ST2 Concentrations With Subclinical Vascular Brain Injury and Incident Stroke. Stroke, 46, 2568–75.

Asundi, J., Zhang, C. L., Donnelly-Roberts, D., Solorio, J. Z., Challagundla, M., Connelly, C., Boch, C., Chen, J., Richter, M., Maneshi, M. M., Swensen, A. M., Lebon, L., Schiffmann, R., Sanyal, S., Sidrauski, C., Kolumam, G. & Baruch, A. 2024. GDF15 is a dynamic biomarker of the integrated stress response in the central nervous system. Cns Neuroscience & Therapeutics, 30.

Baek, S. J., Kim, J. S., Nixon, J. B., Diaugustine, R. P. & Eling, T. E. 2004. Expression of NAG-1, a transforming growth factor-beta superfamily member, by troglitazone requires the early growth response gene EGR-1. J Biol Chem, 279, 6883–92.

Baeva, M. E., Tottenham, I., Koch, M. & Camara-Lemarroy, C. 2023. Biomarkers of disability worsening in inactive primary progressive multiple sclerosis. J Neuroimmunol, 387, 578268.

Ban, N., Siegfried, C. J., Lin, J. B., Shui, Y. B., Sein, J., Pita-Thomas, W., Sene, A., Santeford, A., Gordon, M., Lamb, R., Dong, Z., Kelly, S. C., Cavalli, V., Yoshino, J. & Apte, R. S. 2017. GDF15 is elevated in mice following retinal ganglion cell death and in glaucoma patients. JCI Insight, 2.

Bauskin, A. R., Brown, D. A., Junankar, S., Rasiah, K. K., Eggleton, S., Hunter, M., Liu, T., Smith, D., Kuffner, T., Pankhurst, G. J., Johnen, H., Russell, P. J., Barret, W., Stricker, P. D., Grygiel, J. J., Kench, J. G., Henshall, S. M., Sutherland, R. L. & Breit, S. N. 2005. The Propeptide Mediates Formation of Stromal Stores of PROMIC-1: Role in Determining Prostate Cancer Outcome. Cancer Research, 65, 2330–2336.

Beydoun, M. A., Beydoun, H. A., Noren Hooten, N., Meirelles, O., Li, Z., El-Hajj, Z. W., Weiss, J., Maino Vieytes, C. A., Launer, L. J., Evans, M. K. & Zonderman, A. B. 2023. Hospital-treated prevalent infections, the plasma proteome and incident dementia among UK older adults. iScience, 26, 108526.

Bloch, S. A. A., Lee, J. Y., Syburra, T., Rosendahl, U., Griffiths, M. J. D., Kemp, P. R. & Polkey, M. I. 2015. Increased expression of GDF-15 may mediate ICU-acquired weakness by down-regulating muscle microRNAs. Thorax, 70, 219.

Bonaterra, G. A., Zügel, S., Thogersen, J., Walter, S. A., Haberkorn, U., Strelau, J. & Kinscherf, R. 2012. Growth differentiation factor-15 deficiency inhibits atherosclerosis progression by regulating interleukin-6-dependent inflammatory response to vascular injury. J Am Heart Assoc, 1, e002550.

Bootcov, M. R., Bauskin, A. R., Valenzuela, S. M., Moore, A. G., Bansal, M., He, X. Y., Zhang, H. P., Donnellan, M., Mahler, S., Pryor, K., Walsh, B. J., Nicholson, R. C., Fairlie, W. D., Por, S. B., Robbins, J. M. & Breit, S. N. 1997. MIC-1, a novel macrophage inhibitory cytokine, is a divergent member of the TGF-βlllsuperfamily. Proceedings of the National Academy of Sciences, 94, 11514–11519.

Breit, S. N., Brown, D. A. & Tsai, V. W. 2021. The GDF15-GFRAL Pathway in Health and Metabolic Disease: Friend or Foe? Annu Rev Physiol, 83, 127–151.

Brown, D. A., Hance, K. W., Rogers, C. J., Sansbury, L. B., Albert, P. S., Murphy, G., Laiyemo, A. O., Wang, Z., Cross, A. J., Schatzkin, A., Danta, M., Srasuebkul, P., Amin, J., Law, M., Breit, S. N. & Lanza, E. 2012. Serum macrophage inhibitory cytokine-1 (MIC-1/GDF15): a potential screening tool for the prevention of colon cancer? Cancer Epidemiol Biomarkers Prev, 21, 337–46.

Brown, D. A., Ward, R. L., Buckhaults, P., Liu, T., Romans, K. E., Hawkins, N. J., Bauskin, A. R., Kinzler, K. W., Vogelstein, B. & Breit, S. N. 2003. MIC-1 serum level and genotype: associations with progress and prognosis of colorectal carcinoma. Clin Cancer Res, 9, 2642–50.

Casanova, R., Anderson, A. M., Barnard, R. T., Justice, J. N., Kucharska-Newton, A., Windham, B. G., Palta, P., Gottesman, R. F., Mosley, T. H., Hughes, T. M., Wagenknecht, L. E. & Kritchevsky, S. B. 2023. Is an MRI-derived anatomical measure of dementia risk also a measure of brain aging? GeroScience, 45, 439–450.

Casella, G., Garzetti, L., Gatta, A. T., Finardi, A., Maiorino, C., Ruffini, F., Martino, G., Muzio, L. & Furlan, R. 2016. IL4 induces IL6-producing M2 macrophages associated to inhibition of neuroinflammation in vitro and in vivo. Journal of Neuroinflammation, 13, 139.

Chai, Y. L., Hilal, S., Chong, J. P. C., Ng, Y. X., Liew, O. W., Xu, X., Ikram, M. K., Venketasubramanian, N., Richards, A. M., Lai, M. K. P. & Chen, C. P. 2016. Growth differentiation factor-15 and white matter hyperintensities in cognitive impairment and dementia. Medicine (Baltimore), 95, e4566.

Charalambous, P., Wang, X., Thanos, S., Schober, A. & Unsicker, K. 2013. Regulation and effects of GDF-15 in the retina following optic nerve crush. Cell Tissue Res, 353, 1–8.

Chiariello, A., Valente, S., Pasquinelli, G., Baracca, A., Sgarbi, G., Solaini, G., Medici, V., Fantini, V., Poloni, T. E., Tognocchi, M., Arcaro, M., Galimberti, D., Franceschi, C., Capri, M., Salvioli, S. & Conte, M. 2022. The expression pattern of GDF15 in human brain changes during aging and in Alzheimer’s disease. Front Aging Neurosci, 14, 1058665.

Chung, H. K., Ryu, D., Kim, K. S., Chang, J. Y., Kim, Y. K., Yi, H. S., Kang, S. G., Choi, M. J., Lee, S. E., Jung, S. B., Ryu, M. J., Kim, S. J., Kweon, G. R., Kim, H., Hwang, J. H., Lee, C. H., Lee, S. J., Wall, C. E., Downes, M., Evans, R. M., Auwerx, J. & Shong, M. 2017. Growth differentiation factor 15 is a myomitokine governing systemic energy homeostasis. J Cell Biol, 216, 149–165.

Conte, M., Martucci, M., Mosconi, G., Chiariello, A., Cappuccilli, M., Totti, V., Santoro, A., Franceschi, C. & Salvioli, S. 2020. GDF15 Plasma Level Is Inversely Associated With Level of Physical Activity and Correlates With Markers of Inflammation and Muscle Weakness. Front Immunol, 11, 915.

Conte, M., Ostan, R., Fabbri, C., Santoro, A., Guidarelli, G., Vitale, G., Mari, D., Sevini, F., Capri, M., Sandri, M., Monti, D., Franceschi, C. & Salvioli, S. 2018. Human Aging and Longevity Are Characterized by High Levels of Mitokines. The Journals of Gerontology: Series A, 74, 600–607.

Csecsei, P., Olah, C., Varnai, R., Simon, D., Erdo-Bonyar, S., Berki, T., Czabajszki, M., Zavori, L., Schwarcz, A. & Molnar, T. 2023. Different Kinetics of Serum ADAMTS13, GDF-15, and Neutrophil Gelatinase-Associated Lipocalin in the Early Phase of Aneurysmal Subarachnoid Hemorrhage. Int J Mol Sci, 24.

Danta, M., Barber, D. A., Zhang, H. P., Lee-Ng, M., Baumgart, S. W. L., Tsai, V. W. W., Husaini, Y., Saxena, M., Marquis, C. P., Errington, W., Kerr, S., Breit, S. N. & Brown, D. A. 2017. Macrophage inhibitory cytokine-1/growth differentiation factor-15 as a predictor of colonic neoplasia. Aliment Pharmacol Ther, 46, 347–354.

Davis, R. L., Wong, S. L., Carling, P. J., Payne, T., Sue, C. M. & Bandmann, O. 2020. Serum FGF-21, GDF-15, and blood mtDNA copy number are not biomarkers of Parkinson disease. Neurology-Clinical Practice, 10, 40–46.

De Jager, S. C., Bermúdez, B., Bot, I., Koenen, R. R., Bot, M., Kavelaars, A., De Waard, V., Heijnen, C. J., Muriana, F. J., Weber, C., Van Berkel, T. J., Kuiper, J., Lee, S. J., Abia, R. & Biessen, E. A. 2011. Growth differentiation factor 15 deficiency protects against atherosclerosis by attenuating CCR2-mediated macrophage chemotaxis. J Exp Med, 208, 217–25.

De Marchis, G. M., Krisai, P., Werlen, L., Sinnecker, T., Aeschbacher, S., Dittrich, T. D., Polymeris, A. A., Coslovksy, M., Blum, M. R., Rodondi, N., Reichlin, T., Moschovitis, G., Wuerfel, J., Lyrer, P. A., Fischer, U., Conen, D., Kastner, P., Ziegler, A., Osswald, S., Kuehne, M., Bonati, L. H. & Swiss, A. F. I. 2023. Biomarker, Imaging, and Clinical Factors Associated With Overt and Covert Stroke in Patients With Atrial Fibrillation. Stroke, 54, 2542–2551.

Emmerson, P. J., Wang, F., Du, Y., Liu, Q., Pickard, R. T., Gonciarz, M. D., Coskun, T., Hamang, M. J., Sindelar, D. K., Ballman, K. K., Foltz, L. A., Muppidi, A., Alsina-Fernandez, J., Barnard, G. C., Tang, J. X., Liu, X., Mao, X., Siegel, R., Sloan, J. H., Mitchell, P. J., Zhang, B. B., Gimeno, R. E., Shan, B. & Wu, X. 2017. The metabolic effects of GDF15 are mediated by the orphan receptor GFRAL. Nat Med, 23, 1215–1219.

Evangelisti, S., Gramegna, L. L., La Morgia, C., Di Vito, L., Maresca, A., Talozzi, L., Bianchini, C., Mitolo, M., Manners, D. N., Caporali, L., Valentino, M. L., Liguori, R., Carelli, V., Lodi, R., Testa, C. & Tonon, C. 2022. Molecular biomarkers correlate with brain grey and white matter changes in patients with mitochondrial m.3243A > G mutation. Mol Genet Metab, 135, 72–81.

Fuchs, T., Trollor, J. N., Crawford, J., Brown, D. A., Baune, B. T., Samaras, K., Campbell, L., Breit, S. N., Brodaty, H., Sachdev, P. & Smith, E. 2013. Macrophage inhibitory cytokine-1 is associated with cognitive impairment and predicts cognitive decline - the Sydney Memory and Aging Study. Aging Cell, 12, 882–9.

Golakani, M. H., Mohammad, M. G., Li, H., Gamble, J., Breit, S. N., Ruitenberg, M. J. & Brown, D. A. 2019. MIC-1/GDF15 Overexpression Is Associated with Increased Functional Recovery in Traumatic Spinal Cord Injury. Journal of Neurotrauma, 36, 3410–3421.

Guo, Y., You, J., Zhang, Y., Liu, W. S., Huang, Y. Y., Zhang, Y. R., Zhang, W., Dong, Q., Feng, J. F., Cheng, W. & Yu, J. T. 2024. Plasma proteomic profiles predict future dementia in healthy adults. Nature Aging.

Hsu, J. Y., Crawley, S., Chen, M., Ayupova, D. A., Lindhout, D. A., Higbee, J., Kutach, A., Joo, W., Gao, Z., Fu, D., To, C., Mondal, K., Li, B., Kekatpure, A., Wang, M., Laird, T., Horner, G., Chan, J., Mcentee, M., Lopez, M., Lakshminarasimhan, D., White, A., Wang, S. P., Yao, J., Yie, J., Matern, H., Solloway, M., Haldankar, R., Parsons, T., Tang, J., Shen, W. D., Alice Chen, Y., Tian, H. & Allan, B. B. 2017. Non-homeostatic body weight regulation through a brainstem-restricted receptor for GDF15. Nature, 550, 255–259.

Huddar, A., Govindaraj, P., Chiplunkar, S., Deepha, S., Jessiena Ponmalar, J. N., Philip, M., Nagappa, M., Narayanappa, G., Mahadevan, A., Sinha, S., Taly, A. B. & Parayil Sankaran, B. 2021. Serum fibroblast growth factor 21 and growth differentiation factor 15: Two sensitive biomarkers in the diagnosis of mitochondrial disorders. Mitochondrion, 60, 170–177.

Husaini, Y., Tsai, V. W., Manandhar, R., Zhang, H. P., Lee-Ng, K. K. M., Lebhar, H., Marquis, C. P., Brown, D. A. & Breit, S. N. 2020. Growth differentiation factor-15 slows the growth of murine prostate cancer by stimulating tumor immunity. PLoS One, 15, e0233846.

Iwata, Y., Inagaki, S., Morozumi, W., Nakamura, S., Hara, H. & Shimazawa, M. 2021. Treatment with GDF15, a TGFβ superfamily protein, induces protective effect on retinal ganglion cells. Exp Eye Res, 202, 108338.

Järvilehto, J., Harjuhaahto, S., Palu, E., Auranen, M., Kvist, J., Zetterberg, H., Koskivuori, J., Lehtonen, M., Saukkonen, A. M., Jokela, M., Ylikallio, E. & Tyynismaa, H. 2022. Serum Creatine, Not Neurofilament Light, Is Elevated in CHCHD10-Linked Spinal Muscular Atrophy. Front Neurol, 13, 793937.

Jean, J., Christodoulou, E., Gai, X., Tamrazi, B., Vera, M., Mitchell, W. G. & Schmidt, R. J. 2022. m.3685T > C is a novel mitochondrial DNA variant that causes Leigh syndrome. Cold Spring Harb Mol Case Stud, 8.

Jennings, M. J., Kagiava, A., Vendredy, L., Spaulding, E. L., Stavrou, M., Hathazi, D., Grüneboom, A., De Winter, V., Gess, B., Schara, U., Pogoryelova, O., Lochmüller, H., Borchers, C. H., Roos, A., Burgess, R. W., Timmerman, V., Kleopa, K. A. & Horvath, R. 2022. NCAM1 and GDF15 are biomarkers of Charcot-Marie-Tooth disease in patients and mice. Brain, 145, 3999–4015.

Jeon, H., Kim, H. J., Doo, H. M., Chang, E. H., Kwak, G., Mo, W. M., Jang, S. Y., Lee, M. W., Choi, B. O. & Bin Hong, Y. 2022. Cytokines secreted by mesenchymal stem cells reduce demyelination in an animal model of Charcot-Marie-Tooth disease. Biochemical and Biophysical Research Communications, 597, 1–7.

Jeong, H. S., Shin, J. W., Jeong, J. Y., Kwon, H. J., Koh, H. S., Kim, J. J., Na, K. R., Lee, K. W. & Choi, D. E. 2020. Association of plasma level of growth differentiation factor-15 and clinical outcome after intraarterial thrombectomy. J Stroke Cerebrovasc Dis, 29, 104973.

Ji, F., Chai, Y. L., Liu, S., Kan, C. N., Ong, M., Richards, A. M., Tan, B. Y., Venketasubramanian, N., Pasternak, O., Chen, C., Lai, M. K. P. & Zhou, J. H. 2023. Associations of Blood Cardiovascular Biomarkers With Brain Free Water and Its Relationship to Cognitive Decline: A Diffusion-MRI Study. Neurology, 101, E151–E163.

Ji, X., Zhao, L., Ji, K., Zhao, Y., Li, W., Zhang, R., Hou, Y., Lu, J. & Yan, C. 2017. Growth Differentiation Factor 15 Is a Novel Diagnostic Biomarker of Mitochondrial Diseases. Mol Neurobiol, 54, 8110–8116.

Jiang, J., Trollor, J. N., Brown, D. A., Crawford, J. D., Thalamuthu, A., Smith, E., Breit, S. N., Liu, T., Brodaty, H., Baune, B. T., Sachdev, P. S. & Wen, W. 2015. An inverse relationship between serum macrophage inhibitory cytokine-1 levels and brain white matter integrity in community-dwelling older individuals. Psychoneuroendocrinology, 62, 80–8.

Johnen, H., Lin, S., Kuffner, T., Brown, D. A., Tsai, V. W.-W., Bauskin, A. R., Wu, L., Pankhurst, G., Jiang, L., Junankar, S., Hunter, M., Fairlie, W. D., Lee, N. J., Enriquez, R. F., Baldock, P. A., Corey, E., Apple, F. S., Murakami, M. M., Lin, E.-J., Wang, C., During, M. J., Sainsbury, A., Herzog, H. & Breit, S. N. 2007. Tumor-induced anorexia and weight loss are mediated by the TGF-β superfamily cytokine MIC-1. Nature Medicine, 13, 1333–1340.

Kang, Y. E., Kim, J. M., Lim, M. A., Lee, S. E., Yi, S., Kim, J. T., Oh, C., Liu, L., Jin, Y., Jung, S.-N., Won, H.-R., Chang, J. W., Lee, J. H., Kim, H. J., Koh, H. Y., Jun, S., Cho, S. W., Shong, M. & Koo, B. S. 2020. Growth Differentiation Factor 15 is a Cancer Cell-Induced Mitokine That Primes Thyroid Cancer Cells for Invasiveness. Thyroid®, 31, 772–786.

Karusheva, Y., Ratcliff, M., Mörseburg, A., Barker, P., Melvin, A., Sattar, N., Burling, K., Backmark, A., Roth, R., Jermutus, L., Guiu-Jurado, E., Blüher, M., Welsh, P., Hyvönen, M. & O’rahilly, S. 2022. The Common H202D Variant in GDF-15 Does Not Affect Its Bioactivity but Can Significantly Interfere with Measurement of Its Circulating Levels. J Appl Lab Med, 7, 1388–1400.

Kempf, T., Zarbock, A., Widera, C., Butz, S., Stadtmann, A., Rossaint, J., Bolomini-Vittori, M., Korf-Klingebiel, M., Napp, L. C., Hansen, B., Kanwischer, A., Bavendiek, U., Beutel, G., Hapke, M., Sauer, M. G., Laudanna, C., Hogg, N., Vestweber, D. & Wollert, K. C. 2011. GDF-15 is an inhibitor of leukocyte integrin activation required for survival after myocardial infarction in mice. Nature Medicine, 17, 581–588.

Kim, D. H., Lee, D., Chang, E. H., Kim, J. H., Hwang, J. W., Kim, J. Y., Kyung, J. W., Kim, S. H., Oh, J. S., Shim, S. M., Na, D. L., Oh, W. & Chang, J. W. 2015. GDF-15 secreted from human umbilical cord blood mesenchymal stem cells delivered through the cerebrospinal fluid promotes hippocampal neurogenesis and synaptic activity in an Alzheimer’s disease model. Stem Cells Dev, 24, 2378–90.

Kim, D. H., Lee, D., Lim, H., Choi, S. J., Oh, W., Yang, Y. S., Chang, J. H. & Jeon, H. B. 2018. Effect of growth differentiation factor-15 secreted by human umbilical cord blood-derived mesenchymal stem cells on amyloid beta levels in in vitro and in vivo models of Alzheimer’s disease. Biochem Biophys Res Commun, 504, 933–940.

Klein, A. B., Nicolaisen, T. S., Ørtenblad, N., Gejl, K. D., Jensen, R., Fritzen, A. M., Larsen, E. L., Karstoft, K., Poulsen, H. E., Morville, T., Sahl, R. E., Helge, J. W., Lund, J., Falk, S., Lyngbæk, M., Ellingsgaard, H., Pedersen, B. K., Lu, W., Finan, B., Jørgensen, S. B., Seeley, R. J., Kleinert, M., Kiens, B., Richter, E. A. & Clemmensen, C. 2021. Pharmacological but not physiological GDF15 suppresses feeding and the motivation to exercise. Nat Commun, 12, 1041.

Kosa, P., Wu, T., Phillips, J., Leinonen, M., Masvekar, R., Komori, M., Wichman, A., Sandford, M. & Bielekova, B. 2020. Idebenone does not inhibit disability progression in primary progressive MS. Mult Scler Relat Disord, 45, 102434.

Kostuk, E. W., Cai, J. & Iacovitti, L. 2019. Subregional differences in astrocytes underlie selective neurodegeneration or protection in Parkinson’s disease models in culture. Glia, 67, 1542–1557.

Kuhlmann, T., Moccia, M., Coetzee, T., Cohen, J. A., Correale, J., Graves, J., Marrie, R. A., Montalban, X., Yong, V. W., Thompson, A. J. & Reich, D. S. 2023. Multiple sclerosis progression: time for a new mechanism-driven framework. Lancet Neurol, 22, 78–88.

Kunz, D., Walker, G., Bedoucha, M., Certa, U., März-Weiss, P., Dimitriades-Schmutz, B. & Otten, U. 2009. Expression profiling and Ingenuity biological function analyses of interleukin-6-versus nerve growth factor-stimulated PC12 cells. BMC Genomics, 10, 90.

Lee, L. L., Aung, H. H., Wilson, D. W., Anderson, S. E., Rutledge, J. C. & Rutkowsky, J. M. 2017. Triglyceride-rich lipoprotein lipolysis products increase blood-brain barrier transfer coefficient and induce astrocyte lipid droplets and cell stress. Am J Physiol Cell Physiol, 312, C500–c516.

Li, H., Zhang, G. X., Chen, Y., Xu, H., Fitzgerald, D. C., Zhao, Z. & Rostami, A. 2008. CD11c+CD11b+ dendritic cells play an important role in intravenous tolerance and the suppression of experimental autoimmune encephalomyelitis. J Immunol, 181, 2483–93.

Li, P., Lv, H., Zhang, B., Duan, R., Zhang, X., Lin, P., Song, C. & Liu, Y. 2022a. Growth Differentiation Factor 15 Protects SH-SY5Y Cells From Rotenone-Induced Toxicity by Suppressing Mitochondrial Apoptosis. Front Aging Neurosci, 14, 869558.

Li, P. X., Wong, J., Ayed, A., Ngo, D., Brade, A. M., Arrowsmith, C., Austin, R. C. & Klamut, H. J. 2000. Placental transforming growth factor-beta is a downstream mediator of the growth arrest and apoptotic response of tumor cells to DNA damage and p53 overexpression. J Biol Chem, 275, 20127–35.

Li, Y., Li, S., Qiu, Y., Zhou, M., Chen, M., Hu, Y., Hong, S., Jiang, L. & Guo, Y. 2022b. Circulating FGF21 and GDF15 as Biomarkers for Screening, Diagnosis, and Severity Assessment of Primary Mitochondrial Disorders in Children. Front Pediatr, 10, 851534.

Li, Z., Wang, B., Wu, X., Cheng, S.-Y., Paraoan, L. & Zhou, J. 2005. Identification, expression and functional characterization of the GRAL gene. Journal of Neurochemistry, 95, 361–376.

Lin, J. B., Sheybani, A., Santeford, A. & Apte, R. S. 2021. Longitudinal Growth Differentiation Factor 15 (GDF15) and Long-term Intraocular Pressure Fluctuation in Glaucoma: A Pilot Study. J Ophthalmic Vis Res, 16, 21–27.

Lin, J. B., Sheybani, A., Santeford, A., De Maria, A. & Apte, R. S. 2020. Increased Aqueous Humor GDF15 Is Associated with Worse Visual Field Loss in Pseudoexfoliative Glaucoma Patients. Transl Vis Sci Technol, 9, 16.

Lind, L., Siegbahn, A., Lindahl, B., Stenemo, M., Sundström, J. & Ärnlöv, J. 2015. Discovery of New Risk Markers for Ischemic Stroke Using a Novel Targeted Proteomics Chip. Stroke, 46, 3340–7.

Liu, H., Liu, J., Si, L., Guo, C., Liu, W. & Liu, Y. 2019. GDF-15lllpromoteslllmitochondriallllfunction and proliferation in neuronal HT22 cells. J Cell Biochem, 120, 10530–10547.

Liu, X. 2021. Changes and significance of serum CXCL-16, GDF-15, PLA-2 levels in patients with cerebral infarction. Am J Transl Res, 13, 5617–5622.

Lu, W. H., Giudici, K. V., Morley, J. E., Guyonnet, S., Parini, A., Aggarwal, G., Nguyen, A. D., Li, Y., Bateman, R. J., Vellas, B. & De Souto Barreto, P. 2022. Investigating the combination of plasma amyloid-beta and geroscience biomarkers on the incidence of clinically meaningful cognitive decline in older adults. Geroscience, 44, 1489–1503.

Lu, X., Duan, J., Cheng, Q. & Lu, J. 2020. The association between serum growth differentiation factor-15 and 3-month depression after acute ischemic stroke. J Affect Disord, 260, 695–702.

Machado, V., Gilsbach, R., Das, R., Schober, A., Bogatyreva, L., Hauschke, D., Krieglstein, K., Unsicker, K. & Spittau, B. 2016a. Gdf-15 deficiency does not alter vulnerability of nigrostriatal dopaminergic system in MPTP-intoxicated mice. Cell Tissue Res, 365, 209–23.

Machado, V., Haas, S. J., Von Bohlen Und Halbach, O., Wree, A., Krieglstein, K., Unsicker, K. & Spittau, B. 2016b. Growth/differentiation factor-15 deficiency compromises dopaminergic neuron survival and microglial response in the 6-hydroxydopamine mouse model of Parkinson’s disease. Neurobiol Dis, 88, 1–15.

Mackiewicz, A., Schooltink, H., Heinrich, P. C. & Rose-John, S. 1992. Complex of soluble human IL-6-receptor/IL-6 up-regulates expression of acute-phase proteins. J Immunol, 149, 2021–7.

Maetzler, W., Deleersnijder, W., Hanssens, V., Bernard, A., Brockmann, K., Marquetand, J., Wurster, I., Rattay, T. W., Roncoroni, L., Schaeffer, E., Lerche, S., Apel, A., Deuschle, C. & Berg, D. 2016. GDF15/MIC1 and MMP9 Cerebrospinal Fluid Levels in Parkinson’s Disease and Lewy Body Dementia. Plos One, 11.

Mahato, A. K. & Sidorova, Y. A. 2020. RET Receptor Tyrosine Kinase: Role in Neurodegeneration, Obesity, and Cancer. International Journal of Molecular Sciences, 21, 7108.

Maresca, A., Del Dotto, V., Romagnoli, M., La Morgia, C., Di Vito, L., Capristo, M., Valentino, M. L. & Carelli, V. 2020. Expanding and validating the biomarkers for mitochondrial diseases. J Mol Med (Berl), 98, 1467–1478.

Mcgrath, E. R., Himali, J. J., Levy, D., Conner, S. C., Decarli, C., Pase, M. P., Ninomiya, T., Ohara, T., Courchesne, P., Satizabal, C. L., Vasan, R. S., Beiser, A. S. & Seshadri, S. 2020. Growth Differentiation Factor 15 and NT-proBNP as Blood-Based Markers of Vascular Brain Injury and Dementia. J Am Heart Assoc, 9, e014659.

Melvin, A., Lacerda, E., Dockrell, H. M., O’rahilly, S. & Nacul, L. 2019. Circulating levels of GDF15 in patients with myalgic encephalomyelitis/chronic fatigue syndrome. J Transl Med, 17, 409.

Mendsaikhan, A., Takeuchi, S., Walker, D. G. & Tooyama, I. 2018. Differences in Gene Expression Profiles and Phenotypes of Differentiated SH-SY5Y Neurons Stably Overexpressing Mitochondrial Ferritin. Front Mol Neurosci, 11, 470.

Mensching, L., Börger, A. K., Wang, X., Charalambous, P., Unsicker, K. & Haastert-Talini, K. 2012. Local substitution of GDF-15 improves axonal and sensory recovery after peripheral nerve injury. Cell Tissue Res, 350, 225–38.

Miyaue, N., Yabe, H. & Nagai, M. 2020a. Serum growth differentiation factor 15 levels and clinical manifestations in patients with thiamine deficiency. Neurology and Clinical Neuroscience, 8, 245–250.

Miyaue, N., Yabe, H. & Nagai, M. 2020b. Serum growth differentiation factor 15, but not lactate, is elevated in patients with Parkinson’s disease. J Neurol Sci, 409, 116616.

Montero, R., Yubero, D., Villarroya, J., Henares, D., Jou, C., Rodríguez, M. A., Ramos, F., Nascimento, A., Ortez, C. I., Campistol, J., Perez-Dueñas, B., O’callaghan, M., Pineda, M., Garcia-Cazorla, A., Oferil, J. C., Montoya, J., Ruiz-Pesini, E., Emperador, S., Meznaric, M., Campderros, L., Kalko, S. G., Villarroya, F., Artuch, R. & Jimenez-Mallebrera, C. 2016. GDF-15 Is Elevated in Children with Mitochondrial Diseases and Is Induced by Mitochondrial Dysfunction. PLoS One, 11, e0148709.

Moon, J. S., Goeminne, L. J. E., Kim, J. T., Tian, J. W., Kim, S. H., Nga, H. T., Kang, S. G., Kang, B. E., Byun, J. S., Lee, Y. S., Jeon, J. H., Shong, M., Auwerx, J., Ryu, D. & Yi, H. S. 2020. Growth differentiation factor 15 protects against the aging-mediated systemic inflammatory response in humans and mice. Aging Cell, 19, e13195.

Mullican, S. E., Lin-Schmidt, X., Chin, C. N., Chavez, J. A., Furman, J. L., Armstrong, A. A., Beck, S. C., South, V. J., Dinh, T. Q., Cash-Mason, T. D., Cavanaugh, C. R., Nelson, S., Huang, C., Hunter, M. J. & Rangwala, S. M. 2017. GFRAL is the receptor for GDF15 and the ligand promotes weight loss in mice and nonhuman primates. Nat Med, 23, 1150–1157.

Nichterwitz, S., Nijssen, J., Storvall, H., Schweingruber, C., Comley, L. H., Allodi, I., Lee, M. V., Deng, Q., Sandberg, R. & Hedlund, E. 2020. LCM-seq reveals unique transcriptional adaptation mechanisms of resistant neurons and identifies protective pathways in spinal muscular atrophy. Genome Res, 30, 1083–1096.

Nohara, S., Ishii, A., Yamamoto, F., Yanagiha, K., Moriyama, T., Tozaka, N., Miyake, Z., Yatsuga, S., Koga, Y., Hosaka, T., Terada, M., Yamaguchi, T., Aizawa, S., Mamada, N., Tsuji, H., Tomidokoro, Y., Nakamagoe, K., Ishii, K., Watanabe, M. & Tamaoka, A. 2019. GDF-15, a mitochondrial disease biomarker, is associated with the severity of multiple sclerosis. J Neurol Sci, 405, 116429.

Olsen, O. E., Skjærvik, A., Størdal, B. F., Sundan, A. & Holien, T. 2017. TGF-β contamination of purified recombinant GDF15. PLoS One, 12, e0187349.

Palu, E., Järvilehto, J., Pennonen, J., Huber, N., Herukka, S. K., Haapasalo, A., Isohanni, P., Tyynismaa, H., Auranen, M. & Ylikallio, E. 2023. Rare PMP22 variants in mild to severe neuropathy uncorrelated to plasma GDF15 or neurofilament light. Neurogenetics, 24, 291–301.

Passaro, A. P., Lebos, A. L., Yao, Y. & Stice, S. L. 2021. Immune Response in Neurological Pathology: Emerging Role of Central and Peripheral Immune Crosstalk. Frontiers in Immunology, 12.

Patani, R., Hardingham, G. E. & Liddelow, S. A. 2023. Functional roles of reactive astrocytes in neuroinflammation and neurodegeneration. Nature Reviews Neurology, 19, 395–409.

Patel, S., Alvarez-Guaita, A., Melvin, A., Rimmington, D., Dattilo, A., Miedzybrodzka, E. L., Cimino, I., Maurin, A. C., Roberts, G. P., Meek, C. L., Virtue, S., Sparks, L. M., Parsons, S. A., Redman, L. M., Bray, G. A., Liou, A. P., Woods, R. M., Parry, S. A., Jeppesen, P. B., Kolnes, A. J., Harding, H. P., Ron, D., Vidal-Puig, A., Reimann, F., Gribble, F. M., Hulston, C. J., Farooqi, I. S., Fafournoux, P., Smith, S. R., Jensen, J., Breen, D., Wu, Z., Zhang, B. B., Coll, A. P., Savage, D. B. & O’rahilly, S. 2019. GDF15 Provides an Endocrine Signal of Nutritional Stress in Mice and Humans. Cell Metab, 29, 707–718.e8.

Peñas, A., Fernández-De La Torre, M., Laine-Menéndez, S., Lora, D., Illescas, M., García-Bartolomé, A., Morales-Conejo, M., Arenas, J., Martín, M. A., Morán, M., Domínguez-González, C. & Ugalde, C. 2021. Plasma Gelsolin Reinforces the Diagnostic Value of FGF-21 and GDF-15 for Mitochondrial Disorders. Int J Mol Sci, 22.

Preusch, M. R., Baeuerle, M., Albrecht, C., Blessing, E., Bischof, M., Katus, H. A. & Bea, F. 2013. GDF-15 protects from macrophage accumulation in a mousemodel of advanced atherosclerosis. Eur J Med Res, 18, 19.

Reinehr, S., Safaei, A., Grotegut, P., Guntermann, A., Tsai, T., Hahn, S. A., Kösters, S., Theiss, C., Marcus, K., Dick, H. B., May, C. & Joachim, S. C. 2022. Heat Shock Protein Upregulation Supplemental to Complex mRNA Alterations in Autoimmune Glaucoma. Biomolecules, 12.

Riley, L. G., Nafisinia, M., Menezes, M. J., Nambiar, R., Williams, A., Barnes, E. H., Selvanathan, A., Lichkus, K., Bratkovic, D., Yaplito-Lee, J., Bhattacharya, K., Ellaway, C., Kava, M., Balasubramaniam, S. & Christodoulou, J. 2022. FGF21 outperforms GDF15 as a diagnostic biomarker of mitochondrial disease in children. Mol Genet Metab, 135, 63–71.

Rochette, L., Zeller, M., Cottin, Y. & Vergely, C. 2020. Insights Into Mechanisms of GDF15 and Receptor GFRAL: Therapeutic Targets. Trends Endocrinol Metab, 31, 939–951.

Satoh, J., Tabunoki, H., Nanri, Y., Arima, K. & Yamamura, T. 2006. Human astrocytes express 14-3-3 sigma in response to oxidative and DNA-damaging stresses. Neuroscience Research, 56, 61–72.

Schindowski, K., Von Bohlen Und Halbach, O., Strelau, J., Ridder, D. A., Herrmann, O., Schober, A., Schwaninger, M. & Unsicker, K. 2011. Regulation of GDF-15, a distant TGF-beta superfamily member, in a mouse model of cerebral ischemia. Cell and Tissue Research, 343, 399–409.

Schober, A., Böttner, M., Strelau, J., Kinscherf, R., Bonaterra, G. A., Barth, M., Schilling, L., Fairlie, W. D., Breit, S. N. & Unsicker, K. 2001. Expression of growth differentiation factor-15/ macrophage inhibitory cytokine-1 (GDF-15/MIC-1) in the perinatal, adult, and injured rat brain. J Comp Neurol, 439, 32–45.

Scott, G., Zetterberg, H., Jolly, A., Cole, J. H., De Simoni, S., Jenkins, P. O., Feeney, C., Owen, D. R., Lingford-Hughes, A., Howes, O., Patel, M. C., Goldstone, A. P., Gunn, R. N., Blennow, K., Matthews, P. M. & Sharp, D. J. 2017. Minocycline reduces chronic microglial activation after brain trauma but increases neurodegeneration. Brain, 141, 459–471.

Shvetcov, A., Ruitenberg, M. J., Delerue, F., Gold, W. A., Brown, D. A. & Finney, C. A. 2023. The neuroprotective effects of estrogen and estrogenic compounds in spinal cord injury. Neuroscience & Biobehavioral Reviews, 146, 105074.

Skaper, S. D., Facci, L., Zusso, M. & Giusti, P. 2018. An Inflammation-Centric View of Neurological Disease: Beyond the Neuron. Front Cell Neurosci, 12, 72.

Straub, I. R., Weraarpachai, W. & Shoubridge, E. A. 2021. Multi-OMICS study of a CHCHD10 variant causing ALS demonstrates metabolic rewiring and activation of endoplasmic reticulum and mitochondrial unfolded protein responses. Hum Mol Genet, 30, 687–705.

Strelau, J., Strzelczyk, A., Rusu, P., Bendner, G., Wiese, S., Diella, F., Altick, A. L., Von Bartheld, C. S., Klein, R., Sendtner, M. & Unsicker, K. 2009. Progressive postnatal motoneuron loss in mice lacking GDF-15. J Neurosci, 29, 13640–8.

Strelau, J., Sullivan, A., Böttner, M., Lingor, P., Falkenstein, E., Suter-Crazzolara, C., Galter, D., Jaszai, J., Krieglstein, K. & Unsicker, K. 2000. Growth/differentiation factor-15/macrophage inhibitory cytokine-1 is a novel trophic factor for midbrain dopaminergic neurons in vivo. J Neurosci, 20, 8597–603.

Subramaniam, S., Strelau, J. & Unsicker, K. 2003. Growth differentiation factor-15 prevents low potassium-induced cell death of cerebellar granule neurons by differential regulation of Akt and ERK pathways. J Biol Chem, 278, 8904–12.

Tsai, V. W. W., Husaini, Y., Sainsbury, A., Brown, D. A. & Breit, S. N. 2018. The MIC-1/GDF15-GFRAL Pathway in Energy Homeostasis: Implications for Obesity, Cachexia, and Other Associated Diseases. Cell Metab, 28, 353–368.

Varadarajan, S., Breda, C., Smalley, J. L., Butterworth, M., Farrow, S. N., Giorgini, F. & Cohen, G. M. 2015. The transrepression arm of glucocorticoid receptor signaling is protective in mutant huntingtin-mediated neurodegeneration. Cell Death Differ, 22, 1388–96.

Walker, K. A., Chen, J., Shi, L., Yang, Y., Fornage, M., Zhou, L., Schlosser, P., Surapaneni, A., Grams, M. E., Duggan, M. R., Peng, Z., Gomez, G. T., Tin, A., Hoogeveen, R. C., Sullivan, K. J., Ganz, P., Lindbohm, J. V., Kivimaki, M., Nevado-Holgado, A. J., Buckley, N., Gottesman, R. F., Mosley, T. H., Boerwinkle, E., Ballantyne, C. M. & Coresh, J. 2023. Proteomics analysis of plasma from middle-aged adults identifies protein markers of dementia risk in later life. Science Translational Medicine, 15.

Wang, B., Fu, C., Wei, Y., Xu, B., Yang, R., Li, C., Qiu, M., Yin, Y. & Qin, D. 2022. Ferroptosis-related biomarkers for Alzheimer’s disease: Identification by bioinformatic analysis in hippocampus. Front Cell Neurosci, 16, 1023947.

Wang, X., Krebbers, J., Charalambous, P., Machado, V., Schober, A., Bosse, F., Müller, H. W. & Unsicker, K. 2015. Growth/differentiation factor-15 and its role in peripheral nervous system lesion and regeneration. Cell Tissue Res, 362, 317–30.

Wiklund, F. E., Bennet, A. M., Magnusson, P. K., Eriksson, U. K., Lindmark, F., Wu, L., Yaghoutyfam, N., Marquis, C. P., Stattin, P., Pedersen, N. L., Adami, H. O., Grönberg, H., Breit, S. N. & Brown, D. A. 2010. Macrophage inhibitory cytokine-1 (MIC-1/GDF15): a new marker of all-cause mortality. Aging Cell, 9, 1057–64.

Wollert, K. C., Kempf, T. & Wallentin, L. 2017. Growth Differentiation Factor 15 as a Biomarker in Cardiovascular Disease. Clin Chem, 63, 140–151.

Worthmann, H., Kempf, T., Widera, C., Tryc, A. B., Goldbecker, A., Ma, Y. T., Deb, M., Tountopoulou, A., Lambrecht, J., Heeren, M., Lichtinghagen, R., Wollert, K. C. & Weissenborn, K. 2011. Growth differentiation factor 15 plasma levels and outcome after ischemic stroke. Cerebrovasc Dis, 32, 72–8.

Wu, P. F., Zhang, X. H., Zhou, P., Yin, R., Zhou, X. T. & Zhang, W. 2021. Growth Differentiation Factor 15 Is Associated With Alzheimer’s Disease Risk. Front Genet, 12, 700371.

Xiong, W., Li, D., Feng, Y., Jia, C., Zhang, X. & Liu, Z. 2023. CircLPAR1 Promotes Neuroinflammation and Oxidative Stress in APP/PS1 Mice by Inhibiting SIRT1/Nrf-2/HO-1 Axis Through Destabilizing GDF-15 mRNA. Mol Neurobiol, 60, 2236–2251.

Xiong, W. P., Yao, W. Q., Wang, B. & Liu, K. 2021. BMSCs-exosomes containing GDF-15 alleviated SH-SY5Y cell injury model of Alzheimer’s disease via AKT/GSK-3 beta/beta-catenin. Brain Research Bulletin, 177, 92–102.

Yang, L. D., Chang, C. C., Sun, Z., Madsen, D., Zhu, H. S., Padkjaer, S. B., Wu, X. A., Huang, T., Hultman, K., Paulsen, S. J., Wang, J. S., Bugge, A., Frantzen, J. B., Norgaard, P., Jeppesen, J. F., Yang, Z. R., Secher, A., Chen, H. B., Li, X., John, L. M., Shan, B., He, Z. H., Gao, X., Su, J., Hansen, K. T., Yang, W. & Jorgensen, S. B. 2017. GFRAL is the receptor for GDF15 and is required for the anti-obesity effects of the ligand. Nature Medicine, 23, 1158-+.

Yao, X., Wang, D., Zhang, L., Wang, L., Zhao, Z., Chen, S., Wang, X., Yue, T. & Liu, Y. 2017. Serum Growth Differentiation Factor 15 in Parkinson Disease. Neurodegener Dis, 17, 251–260.

Yatsuga, S., Fujita, Y., Ishii, A., Fukumoto, Y., Arahata, H., Kakuma, T., Kojima, T., Ito, M., Tanaka, M., Saiki, R. & Koga, Y. 2015. Growth differentiation factor 15 as a useful biomarker for mitochondrial disorders. Ann Neurol, 78, 814–23.

Yi, M. H., Zhang, E., Baek, H., Kim, S., Shin, N., Kang, J. W., Lee, S., Oh, S. H. & Kim, D. W. 2015. Growth Differentiation Factor 15 Expression in Astrocytes After Excitotoxic Lesion in the Mouse Hippocampus. Exp Neurobiol, 24, 133–8.

You, J., Guo, Y., Zhang, Y., Kang, J. J., Wang, L. B., Feng, J. F., Cheng, W. & Yu, J. T. 2023. Plasma proteomic profiles predict individual future health risk. Nat Commun, 14, 7817.

Younes, R., Issa, Y., Jdaa, N., Chouaib, B., Brugioti, V., Challuau, D., Raoul, C., Scamps, F., Cuisinier, F. & Hilaire, C. 2023. The Secretome of Human Dental Pulp Stem Cells and Its Components GDF15 and HB-EGF Protect Amyotrophic Lateral Sclerosis Motoneurons against Death. Biomedicines, 11.

Yue, T., Lu, H., Yao, X. M., Du, X., Wang, L. L., Guo, D. D. & Liu, Y. M. 2020. Elevated serum growth differentiation factor 15 in multiple system atrophy patients: A case control study. World J Clin Cases, 8, 2473–2483.

Zhang, B., Chang, J. Y., Lee, M. H., Ju, S. H., Yi, H. S. & Shong, M. 2024. Mitochondrial Stress and Mitokines: Therapeutic Perspectives for the Treatment of Metabolic Diseases. Diabetes Metab J.

Zhang, Y., Zhan, Y., Kou, Y., Yin, X., Wang, Y. & Zhang, D. 2020. Identification of biological pathways and genes associated with neurogenic heterotopic ossification by text mining. PeerJ, 8, e8276.

